# Rapid evolution of mouse saliva through regulatory rewiring and gene expansions

**DOI:** 10.64898/2026.03.18.712472

**Authors:** Luane Jandira Bueno Landau, Shikha Jain, Nathan Griffin, Achisha Saikia, Jill M. Kramer, Sarah Knox, Stefan Ruhl, Omer Gokcumen

## Abstract

**Background:** Mammalian saliva plays essential roles in digestion, immunity, and host-microbiome interactions, yet its protein composition varies widely across species and sexes. This diversity makes saliva a powerful model to study evolution of gene expression. Here, we compared mouse and human salivary glands at the genomic, transcriptomic, and proteomic levels to understand how saliva composition evolves.

**Results:** We found that evolution of gene expression in mouse salivary glands is driven by rapid gene turnover and sexual dimorphism. In the submandibular and sublingual glands, 68% and 73% of expression from genes encoding secreted proteins derives from lineage-specific genes that lack one-to-one human orthologs. This contrasts the prevailing view that gene expression is largely conserved across tissues and highlights saliva as an unusually dynamic system of molecular evolution. Mouse submandibular gland shows striking sexual dimorphism, with 1537 tissue specific sex-biased genes, 3.8 times higher than in the liver (p<0.001), a classic model of sex-biased expression. We also identified sex-specific expression of glycosylation genes associated with differential Muc19 sialylation. These results suggest that sexual dimorphism in mouse saliva arises from both transcriptional differences in secreted proteins and sex-specific post-translational modifications. Further, differentially expressed genes in submandibular gland cluster in genomic regions shaped by recent gene duplication, such as the kallikrein gene cluster, a mouse-specific expansion that accounts for ∼16.4% of male-biased submandibular gland expression. Our analyses suggest that this bias arises through regulatory changes that are expanded by gene duplication, including the spread of a testosterone-associated regulatory motif and the expansion of a shared chromatin domain that promotes coordinated gene regulation.

**Conclusions:** Together, our results reveal that lineage-specific genes disproportionately shape the mouse salivary transcriptome and that gene duplication and regulatory rewiring contribute to the rapid evolution of sex-biased expression in the submandibular gland. We describe a novel mechanism through which lineage-specific gene duplication and regulatory rewiring drive rapid, sex-specific evolution of the mammalian saliva. Our findings support saliva as an evolutionarily dynamic system and are consistent with developmental systems drift, whereby similar phenotypes across species may be maintained despite substantial divergence in their underlying gene-regulatory programs.

## INTRODUCTION

Diet and pathogen exposure are major environmental drivers of mammalian evolution [1, 2]. Because the oral cavity represents the body’s first interface with the external environment, saliva plays a central role in mediating these pressures, balancing protection against pathogens with responsiveness to dietary variation. Saliva also contributes to oral homeostasis by lubricating and cleansing the mouth, protecting tooth enamel, enabling taste perception, and initiating digestion [3, 4]. These functions are carried out by a complex mixture of proteins secreted primarily by the three major salivary glands, parotid, submandibular, and sublingual glands[5, 6]. Importantly, the salivary gland secretome, i.e., the entirety of proteins secreted by the salivary glands, reflects adaptive tuning to species-specific ecological contexts, with protein repertoires shaped by dietary specialization [7] and pathogen exposure [8]. Together, the integration of immune, digestive, and sensory functions makes saliva an exceptional system for investigating how environmental pressures drive the evolution of mammalian genomes.

Previous studies suggested that saliva-related genes undergo rapid evolution, mainly to accommodate environmental and dietary changes which accompany speciation [7]. Recent locus-specific comparative genomic studies showed that genes expressed in the salivary glands that encode secreted salivary proteins differ substantially between mice and humans [9]. For instance, analyses of the secretory calcium-binding phosphoprotein (SCPP) locus indicated that multiple human salivary genes lack orthologs in mice. Proteomic analyses also have revealed that the composition of saliva proteins differs markedly between humans and mice [10, 11], and comparative studies further show substantial interspecies differences in their major components [12]. These findings suggest that human and mouse salivary glands differ greatly in both gene content and expression profiles, supporting the view that the salivary gland secretome evolves rapidly in response to environmental and pathogenic pressures. Despite these insights, a systematic, secretome-wide investigation of the evolutionary dynamics and regulatory mechanisms underlying these differences has not yet been conducted.

In contrast to humans, the salivary glands in many other mammalian species are different between sexes. Sexual dimorphism in salivary glands has been well documented in rodents [13], and pronounced sex differences have been shown in salivary glands of mice [14], which are not known to occur in humans. Generally, sex-biased genes across multiple tissues in mammals are poorly conserved across species [15]. In particular, the mouse submandibular gland exhibits striking sex-specific differences in cellular composition, secretory activity, and hormonal regulation [16–18]. Although substantial progress has been made in understanding saliva composition and its role in oral biology in both species, the evolutionary basis of lineage-and sex-specific variation in the salivary gland secretome remains underexplored.

Here, we investigate the mechanisms underlying the rapid diversification of the salivary gland secretome. Using comparative analyses between humans and mice, we assess the conservation of genes highly expressed in salivary glands and determine the contribution of lineage-specific genes to the salivary gland secretome. We further compare male and female mice to identify the genomic basis of sex-biased expression in the submandibular gland. By integrating transcriptomic, proteomic, and genomic data, we delineate the evolutionary processes driving salivary diversity and sexual dimorphism.

## RESULTS

### The highest proportion of gene expression in mouse submandibular and sublingual glands derives from lineage-specific genes

To investigate differences in salivary gene expression across commonly used mouse models and compare them to humans, we performed transcriptomic profiling of the parotid, submandibular, and sublingual glands, along with the pancreas and liver as reference tissues, in CD1 and C57BL/6 strains (n = 3 per sex per strain; **Tables S1**). Both CD1 and C57BL/6 strains are widely used in laboratory studies of salivary glands, and were included as a representative of an inbred and outbred mouse strain, respectively, to strengthen the broader applicability of our findings [19–21]. In parallel, we conducted mass spectrometry-based proteomic profiling of whole saliva from ten C57BL/6 mice, five males and five females (**Table S2**). Principal component analysis (PCA) of the transcriptomic data shows that samples cluster strongly by tissue of origin, with clear separation among the three salivary glands (**Figure S1**), supporting the robustness of the dataset.

To characterize gland-specific expression patterns and contribution to saliva proteome, we calculated mean transcript abundance herein estimated in transcripts per million (TPM). Each gland displayed a distinct transcriptional signature, typically dominated by a small subset of highly expressed, gland-specific genes that code for secreted proteins, consistent with previous work [22]. In this sense, salivary glands function as secretory “pumps”, where a small number of genes encoding abundant, secreted proteins account for a large share of total transcriptional output: fewer than 10 genes collectively account for more than 50% of total expression in mouse parotid and submandibular glands. A similarly skewed distribution is seen in humans, where fewer than five genes account for more than 50% of total expression in each salivary gland, compared with 128 genes in the liver. Thus, although many genes are detectably expressed in the salivary glands, a small subset contributes disproportionately to total transcript abundance.

To further investigate these highly expressed genes, we identified the 30 most highly expressed genes encoding secreted proteins in each mouse gland (see **Methods**; **Figure 1a, Table S3**). These genes account for 69.3%, 64.9%, and 35.6% of total transcript abundance in the parotid, submandibular, and sublingual glands, respectively. When considering only transcripts encoding secreted proteins, these top 30 genes explain an even larger fraction of expression: 95.7%, 95.9%, and 87.0% in the respective glands. To further confirm that these results were robust to the choice of normalization and mapping approach, we reproduced the analysis using Salmon [23], a k-mer based mapping algorithm (see **Methods**). Using this approach, we similarly found that, across the salivary glands, the top 30 genes encoding secreted proteins accounted for more than 50% of total expression and more than 90% of the expression attributable to secreted proteins (**Table S4**). These findings support the robustness of our results across analytical approaches and suggest that the observed pattern likely reflects a biological effect consistent with the “pump” model described above.

**Figure 1.**
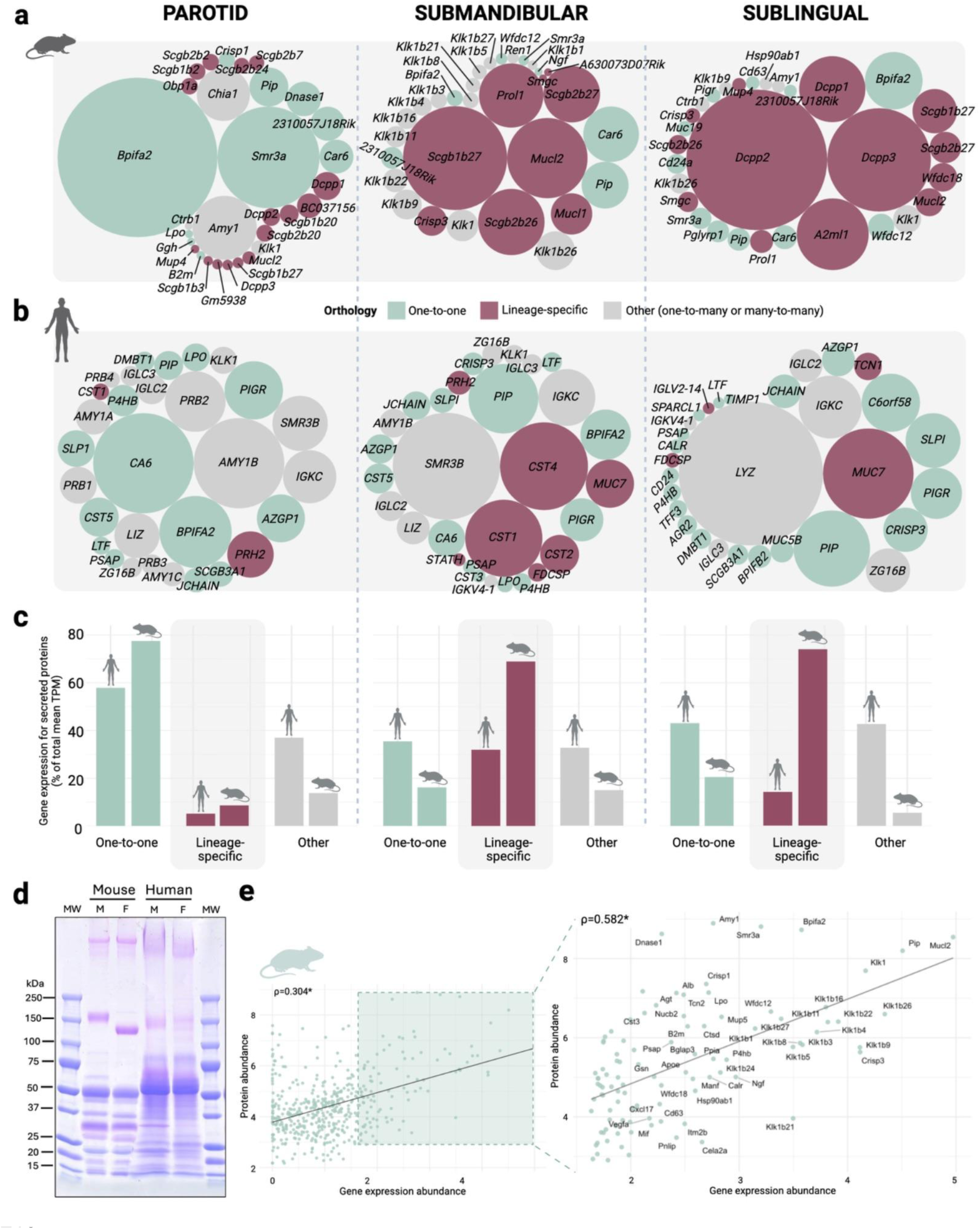
Most mouse genes coding for secreted proteins in salivary glands are lineage-specific. **(a)** Bubble plots showing the top 30 most highly transcribed genes that code for secreted proteins in each major mouse salivary gland. Bubble size represents mean transcript abundance herein estimated in transcripts per million (TPM), and colors denote the type of orthology: one-to-one (green), lineage-specific (maroon), and other orthologous relationships (including many-to-many and one-to-many) (gray). **(b)** Same analysis as in A, here for human salivary glands. (**c**) Proportion of total gene expression (mean TPM) of genes that code for secreted proteins attributed to each orthology category in mouse and human. **(d)** Saliva proteins separated by SDS-PAGE and stained with Coomassie blue and Periodic acid Schiff’s stain for pooled whole saliva of males (M, n=5) and females (F, n=5) for mice and humans. Pink bands are glycosylated proteins revealed by periodic acid Schiff stain. Note the shift in gelelectrophoretic mobility of the high-molecular-weight glycoprotein band from 150 kDa in males to 130 kDa in females. MW denotes molecular weight markers. **(e)** Correlation between gene expression (Log10(1+Mean TPM)) and protein abundance (Log10(1+Mean iBAQ)) for genes that code for secreted proteins in the submandibular gland. The right panel highlights the top 100 most highly expressed genes, for which RNA and protein expression are more concordant. ρ denotes Spearman’s correlation. *denotes *p*<0.05.

Next, we categorized mouse salivary gland genes based on their evolutionary relationships to human genes: one-to-one orthologs, other orthologous relationships (i.e., one-to-many or many-to-many, based on the Jackson Laboratory Mouse Genome Informatics database), or lineage-specific genes (**Fig. 1a**). We applied the same orthology classification to human salivary gland transcriptomes from our previous study [24], enabling direct comparison between species (**Figure 1b, Table S5**). The proportion of expression originating from one-to-one orthologs that code for secreted proteins are remarkably low for sublingual and submandibular glands, approximately 8% and 11%, respectively (**Figure 1c**). In contrast, the parotid gland displays a higher proportion of expression of one-to-one orthologs (77.58%), suggesting greater evolutionary conservation in its core secretory functions. To ensure that our choice of orthology database did not bias the results, we repeated this analysis using the Ensembl orthology database and obtained similar results (**Fig. S2, see Methods**).

We recently hypothesized, based on a locus-specific analysis, that salivary glands evolve faster than other tissues because of shifting dietary and pathogenic evolutionary pressures [9]. Here, we tested this hypothesis more comprehensively at the transcriptome-wide level for genes that code for secreted proteins by comparing the expression levels of one-to-one orthologs between human and mouse, which are expected to maintain comparable expression levels across species [25]. We found that Spearman correlations between mouse and human were significantly higher in liver and pancreas (liver: ρ=0.78; pancreas: ρ=0.642) compared to each of the salivary glands (parotid: ρ = 0.575; submandibular: ρ = 0.58; sublingual: ρ = 0.606; p < 0.001 for all comparisons) with the exception of sublingual, that shows no difference with pancreas (*p*_adj_=0.13) (**Fig. S3** and **Fig S4**, **Table S6**). Of note, such cross-species transcriptomic analyses are subject to potential confounders, including batch effects, normalization biases, and annotation differences. Thus, although our results show robust and consistent trends, further verification is warranted.

To determine whether these transcriptomic differences for genes encoding secreted proteins are also reflected at the proteomic level, we performed gel electrophoresis of whole saliva (**Figure 1d**; **Figure S5**). The resulting protein and glycoprotein profiles revealed striking qualitative and quantitative differences in banding patterns between human and mouse saliva, with only a few bands appearing to match between the two species. Particularly noteworthy is the observed shift in gel electrophoretic mobility of the high-molecular-weight glycoprotein band from 150 kDa in males to 130 kDa in females. This protein band was identified by mass spectrometry (MS/MS) analysis as Muc19 (**Table S7**).

In parallel, we asked the extent to which the transcriptomic variation observed translates into variation in the salivary proteome. To assess this, we compared gene expression in the salivary glands of each mouse strain with the abundance of the corresponding proteins in their saliva for 467 gene-protein pairs (see **Methods**). We observed a significant positive correlation for each gland (parotid: ρ = 0.362, p = 7.29 × 10^-16^; sublingual: ρ = 0.268, p = 3.76 × 10^-9^; submandibular: ρ = 0.304, p = 1.81 × 10^-11^) (**Figure S6**). Notably, correlation strength increased among the most highly expressed genes that code for secreted proteins, with stronger associations observed when restricting the analysis to the top 100 transcripts per gland (parotid: ρ = 0.492, p = 2.06×10^-7^; sublingual: ρ = 0.528, p = 1.65×10^-8^; submandibular: ρ = 0.582, p = 2.10×10^-10^) (**Figure 1e**). These protein-level analyses support the notion that transcript abundance in the salivary glands directly affects protein levels in saliva, and that highly expressed genes encoding secreted proteins have an outsized influence on shaping the oral proteome.

Our combined transcriptomic and proteomic analyses support a model in which mammalian salivary glands function as specialized organs, biological pumps, that produce a limited set of highly abundant secreted proteins. Our data further support the hypothesis that the genes coding for secreted proteins in the salivary glands show faster turnover than those in liver and pancreas, a phenomenon more pronounced among the top expressed genes encoding secreted proteins. Transcriptomic data from individual salivary glands accurately predict their relative contributions to the oral proteome, consistent with previous observations in humans [24]. Overall, our results reveal fundamental differences in expression profiles of genes coding for secreted proteins in the salivary glands of humans and mice, consistent with a rapid evolution of the salivary gland secretome.

### The submandibular gland is the major contributor to sexual dimorphism in saliva

Previous studies have documented sexual dimorphism in the postnatal development and cellular composition of the mouse submandibular gland, most notably the male-specific expansion of granular convoluted tubules (ducts) [26, 27]. These morphological differences are known to influence salivary protein output and gene expression [18, 28]. Thus, we hypothesize that the submandibular gland contributes to sexual dimorphism in the salivary proteome. Analysis of our mass spectrometry dataset identified 368 and 152 proteins with significantly male-and female-biased protein abundances, respectively (*p*_adj_<0.05 and |log_2_FoldChange| > 0.5, **Table S8**). To determine the transcriptional basis of this protein-level dimorphism, we next examined whether sex-biased gene expression across individual salivary glands contributes to the observed differences in salivary protein abundance.

To systematically address this hypothesis, we performed comparative transcriptomic analyses of sex-biased gene expression across mouse salivary glands, liver, and pancreas (**Table S9**). Our experimental design uses both strains to robustly identify sex-biased genes (see **Methods**). The submandibular gland exhibited the largest number of tissue-specific differentially expressed genes (DEGs, FoldEnrichment>1, *p_adj_*<0.05) between sexes (1,537 genes) with other salivary glands exhibiting significantly fewer tissue-specific DEGs (**Figure 2a**, p_adj_< 0.001; **Table S10**), consistent with previous reports [18]. The proportion of DEGs in the submandibular gland was approximately 3.8-and 52.8-fold higher than in the liver and pancreas, respectively (p_adj_ < 0.001). What distinguishes our analysis is the direct cross-organ comparison. These results place the submandibular gland among the most sexually dimorphic tissues in mice, clearly exceeding even the liver, which is widely regarded as the canonical model for studying sexual dimorphism in mammalian gene expression [29, 30]. Further, a direct comparison of the extent of sexual differentiation in submandibular transcriptomes shows a dramatic difference between mice and humans (**Figure 2b, Table S11**).

**Figure 2.**
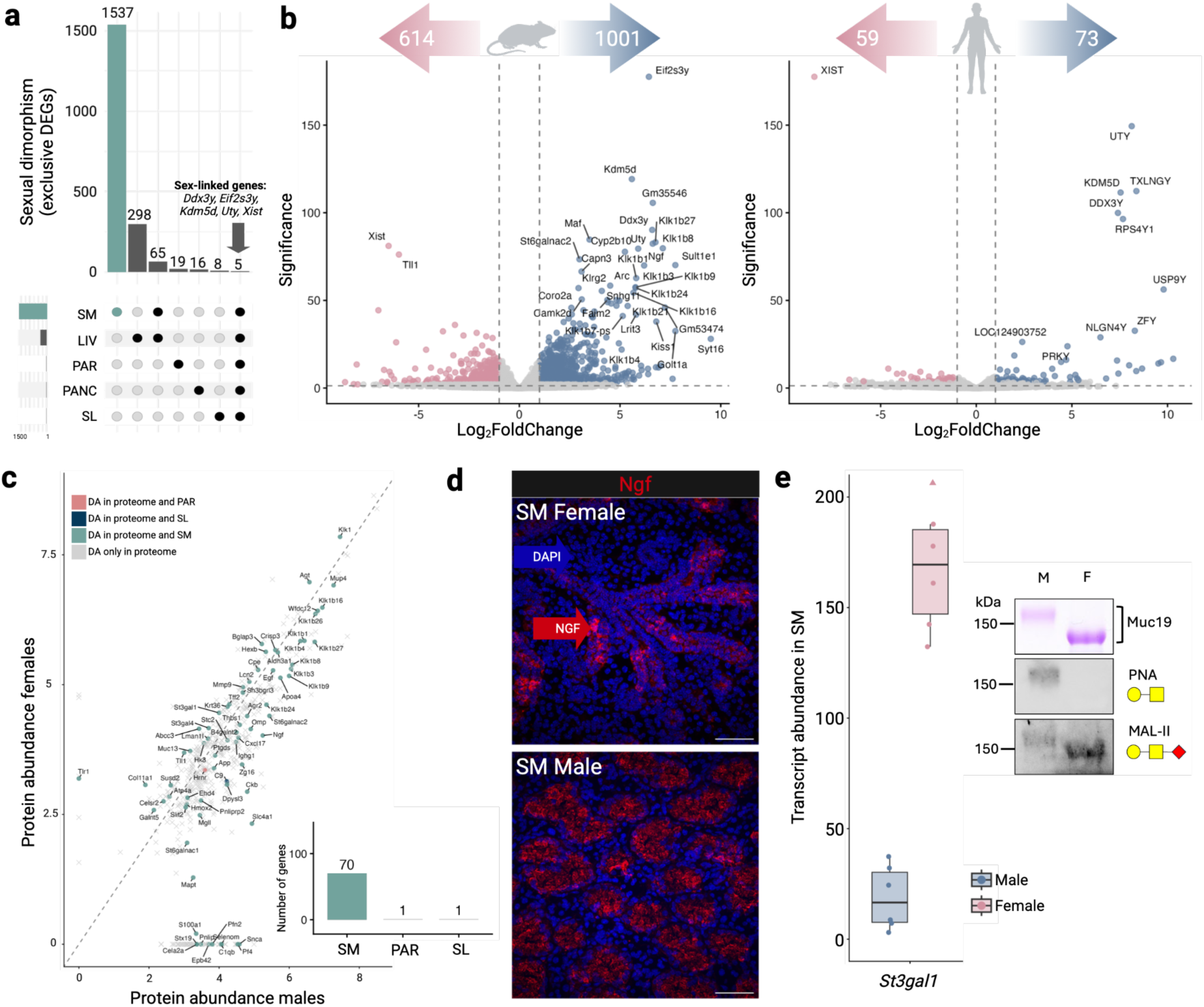
The submandibular gland is the major contributor to sexual dimorphism of the salivary gland secretome. **(a)** Upset plot showing that the submandibular (SM) gland harbors more exclusive DEGs between mouse sexes than other major salivary glands (parotid [PAR], sublingual [SL]) or non-salivary tissues (liver [LIV], pancreas [PANC]). Sex-linked genes (*Ddx3y*, *Eif2s3y*, *Kdm5d*, *Uty*, *Xist*) that are differentially expressed in all tissues are indicated with an arrow. **(b)** Volcano plot of differentially expressed genes in the mouse (left) and human (right) submandibular gland. Arrows indicate the total number of significantly upregulated genes in females (pink) and males (blue). Pink dots represent genes significantly upregulated in females, and blue dots represent genes significantly upregulated in males. Significance is measured in Log10(padj). **(c)**. Correlation plot of protein abundances in males and females quantified in Log10(1+Mean iBAQ). Each dot represents a differentially abundant protein. Female protein abundance is shown on the y-axis and male protein abundance on the x-axis. The dashed line indicates one-to-one correspondence. Colored points denote sexually dimorphic proteins that are also differentially expressed genes in each gland. The inset panel shows the number of differentially abundant proteins whose genes are also differentially expressed in each gland, with the highest contribution observed in the submandibular gland. **(d)** Immunofluorescence staining showing differences in expression patterns between female (top) and male (bottom) submandibular glands. NGF (red). DAPI marks nuclei (blue). **(e)** Sex-biased expression of the sialyltransferase *St3gal1* is associated with differences in Muc19 glycosylation. Left: Boxplot showing transcript abundance of *St3gal1* (TPM) in male and female mice. One female outlier (TPM = 443) is shown as a triangle and was kept for statistical calculations. Right: SDS-PAGE and lectin blot analysis of Muc19 in pooled male (M) and female (F) saliva samples. PNA detects unsialylated core O-glycans, whereas MAL-II detects terminal α2,3-linked sialic acids. Differences in lectin binding and electrophoretic mobility indicate higher sialylation of female Muc19, consistent with increased *St3gal1* expression.

Next, we tested if most sexually dimorphic saliva proteins originate from genes with sex-biased expression in the submandibular gland. Protein abundance in saliva is influenced by many factors, including RNA abundance, translational efficiency, posttranslational modifications, and peptide degradation. Despite this complexity, we found that 70 of the 507 sexually dimorphic salivary proteins are encoded by genes that are differentially expressed in the submandibular gland, compared with only one gene in the parotid gland and one in the sublingual gland (**Figure 2c**). These results show that the submandibular gland is the primary contributor to the sexual dimorphism observed in the protein composition of mouse saliva.

To further validate these results, we used immunohistochemical staining with well-characterized antibodies to examine the expression of nerve growth factor (*Ngf*), one representative sex-biased protein, which showed pronounced sexual dimorphism in both submandibular RNA-seq and salivary proteome analyses [31, 32]. Examination of submandibular gland sections from male and female mice revealed clear immunohistochemical differences between sexes (**Figure 2d**), supporting that pronounced sexual dimorphism at the gene expression and secreted protein levels are also manifested at the cellular level.

In addition to transcriptional differences, post-translational modifications, such as glycosylation, may further contribute to sex-specific differences in the biochemical and functional properties of salivary proteins. In particular, sialic acids are negatively charged terminal glycan residues on most mammalian glycan chain and are commonly present on glycans of secreted glycoproteins, most prominently on mucins, where they influence the biophysical and tribological properties of these molecules and play important roles in their interaction with microbes and viruses [33, 34]. Consistent with this possibility, we observed differential expression of several genes involved in sialylation pathways, including the sialyltransferases *St3gal1*, *St3gal4*, *St3gal5*, *St6galnac1*, and *St6galnac2*, between males and females (**Figure S7**). Notably, in contrast to mice, in humans, none of these sialyltransferases are differentially expressed between sexes in the submandibular gland.

To test whether these transcriptional differences correspond to differences in protein glycosylation, we examined a prominent glycoprotein band in mouse saliva whose electrophoretic mobility shifts from ∼150 kDa in males to ∼130 kDa in females (**Fig. 1d**). Mass spectrometry (MS/MS) analysis of the major protein excised from these bands identified it as the mucin Muc19, a highly O-glycosylated, gel-forming salivary mucin involved in saliva lubrication, hydration, and innate protection of the oral cavity against bacterial colonization [35]. In agreement with previous work [36], we found that Muc19 was highly expressed in the sublingual gland of both sexes (mean TPM=1524.01). Notably, in humans MUC19 is expressed in salivary glands at much lower levels (mean TPM 213.15 in SL), consistent with disputed activity of MUC19 in human saliva [37]. Regardless, our results indicate that the sex bias in this protein’s molecular weight reflects differences in protein size or post-translational modification rather than in protein abundance.

Lectin blotting using Peanut Agglutinin (PNA), revealed strong binding to the band in male saliva, but lack of binding to the band in females, whereas lectin blotting with Maackia amurensis lectin II (MAL-II) showed strong binding to the band in females and weak binding to the band in male saliva. Together, these results indicate presence of unsialylated core O-glycans (Galꞵ1-3GalNAc) on male Muc19 and a higher proportion of terminal α2,3-linked sialylation on female Muc19 (**Figure 2e**). The higher degree of Muc19 sialylation in female mice is consistent with the observed electrophoretic shift of the Muc19 band from ∼150 kDa in male saliva to ∼130 kDa in female saliva due to the added electronegativity conferred by the higher number of negatively charged sialic acid molecules on the female mucin glycan chains. These results suggest that, in addition to sex-specific differences affecting the transcription of genes encoding secreted proteins in the submandibular gland directly, sex-specific post-translational modifications also contribute substantially to the sexual dimorphism observed in mouse saliva, and are caused by sex-specific transcription differences in modifying enzymes, thereby affecting secreted protein products indirectly in a sex-specific manner.

Together, these findings demonstrate that the submandibular gland, by far the most sexually dimorphic tissue in our dataset, is the primary contributor to the strong sexual dimorphism observed in the salivary proteome, with additional contributions from sex-specific post-translational modifications.

### *Klk* gene family expansion underlies sex-biased expression in mice

The profound sexual dimorphism in the mouse submandibular gland is not recapitulated in humans (**Figure 2b**). This observation motivates a closer examination of the evolutionary processes shaping sex-biased gene expression in the mouse submandibular gland. In particular, we asked whether this extensive sex bias might reflect coordinated regulation of clusters of genes within specific genomic regions or a broad, genome-wide effect.

To address this, we first searched for physically clustered genes that may be co-regulated to express in a sex-specific manner. Specifically, we performed a genomic clustering analysis of sex-biased DEGs within 100-kb of each other in the mouse submandibular gland (see **Methods**; **Table S12**). We found that the Klk gene family forms the largest cluster of sex-biased genes in the genome and is a significant outlier relative to other clusters (*p* < 0.001; **Figure 3a**). Fourteen members of the kallikrein (Klk) family (*Klk15, Klk1b8, Klk1b1, Klk1b9, Klk1b11, Klk1b26, Klk1b27, Klk1b21, Klk1b22, Klk1b16, Klk1b24, Klk1b3, Klk1b4, and Klk1b5*) exhibit strong male-biased expression, whereas one member (*Klk1*) shows female-biased expression. Altogether, *Klk* genes make up ∼16.4% of total gene expression in male submandibular glands, compared with only 2.5% in female ones (**Figure 3b**).

**Figure 3.**
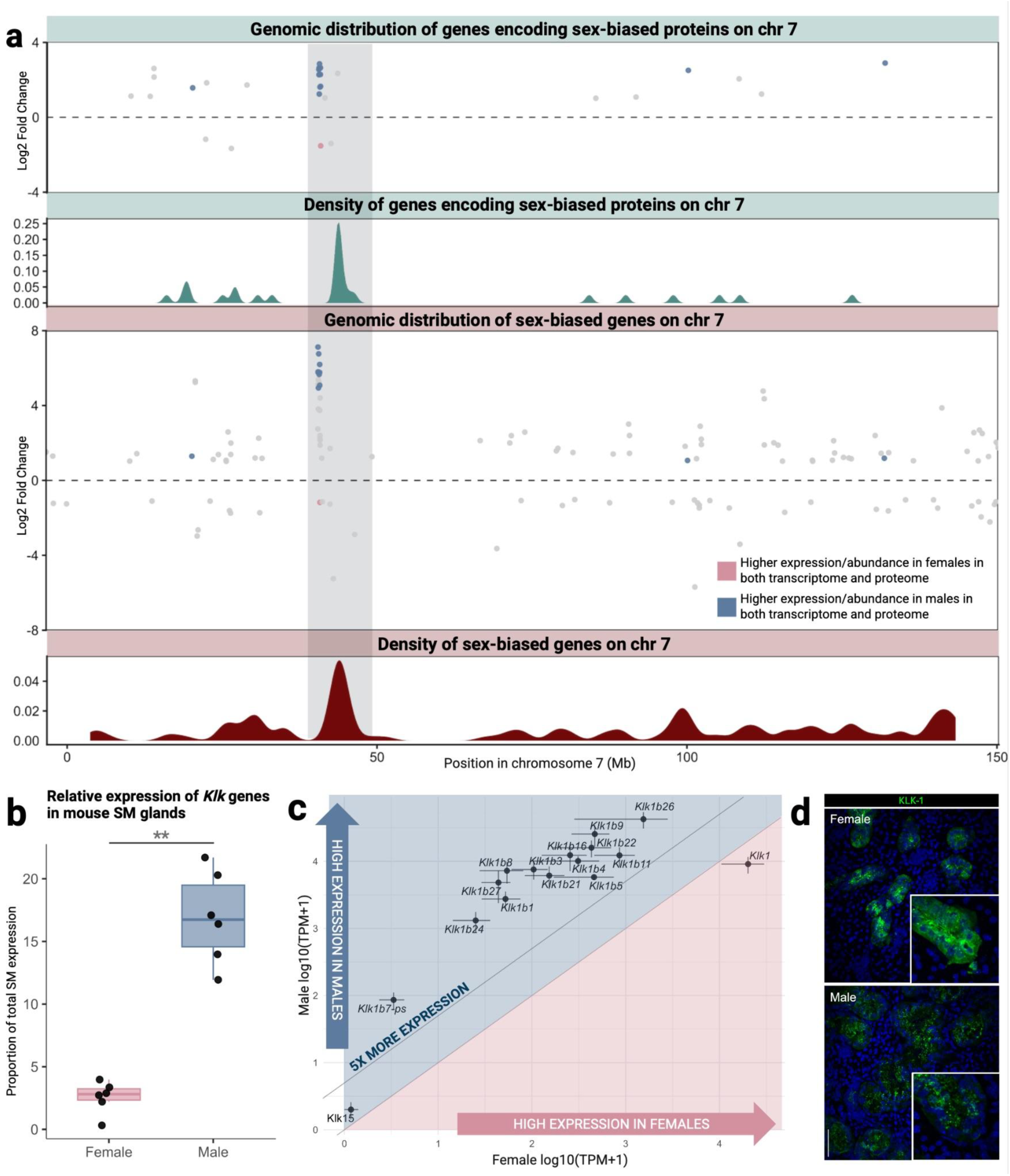
The *Klk* gene family forms the largest cluster of sex-biased transcripts and proteins in mouse submandibular glands, independent of differences in cell abundance. **(a)** Genomic distribution of genes encoding for sex-biased proteins (top, green) and sex-biased transcripts (bottom, magenta) along mouse chromosome 7 in the submandibular gland. Points represent differentially abundant proteins or differentially expressed genes, plotted by genomic position and log2 fold difference between sexes. The shaded region highlights the *Klk* locus. Density tracks below each scatter plot show kernel-smoothed spatial density of significant genes or proteins along the chromosome 7. Blue and pink points indicate genes whose sex bias at the transcript level is also supported by proteomic differences in males and females, respectively. **(b)** Relative expression of *Klk* genes in male and female submandibular glands. Boxes show distribution across biological replicates; points represent individual samples. **\*\*** indicates *p* < 0.001. **(c)** Comparison of female versus male *Klk* gene expression levels. All *Klk1b* paralogs show >5-fold higher expression in males, indicating that sex-biased expression is not explained by differences in glandular cell composition. **(d)** Immunofluorescence staining of submandibular glands from female (top) and male (bottom). Klk1 protein is shown in green; nuclei are counterstained with DAPI (blue), illustrating stronger Klk1 expression in females.

To confirm that the observed sex-biased expression of the *Klk* gene family was not driven by normalization artifacts or mapping differences, we independently validated these results using an alternative Salmon-based pipeline (see **Methods**). This analysis reproduced the sex-biased expression patterns for all of these genes except *Klk1*, which was not significantly different between males and females (**Table S13**). Together, these results support the interpretation that the observed sex bias reflects a biological effect rather than a pipeline-specific artifact. Of note, although some Klk genes are also abundantly expressed in other salivary glands, they do not show the dramatic sex-biased expression observed in the submandibular gland, indicating that the sex bias is tissue-specific. Whether other gene family expansions also contribute to the sex bias observed in submandibular glands remains an important area for future investigation.

We then asked whether the clustering of sex-biased genes can also be observed at the proteomic level. Of the Klk genes showing sex-biased expression, 11 out of 15 also display concordant sex-biased differences in protein abundance in saliva. Using genomic coordinates of genes encoding differentially abundant proteins, we observed that the *Klk* genes exhibiting sex-biased protein abundance are clustered on chromosome 7 (**Figure 3a**), consistent with the transcriptomic clustering pattern. Together, these results indicate that the kallikrein locus represents a genome-wide outlier cluster of coordinated, sex-biased regulation at both the transcriptomic and proteomic levels.

Previous studies have already shown that *Klk* genes exhibit male-biased expression in the mouse submandibular gland [18, 38] and in saliva [10, 39–41]. Single-cell transcriptomic analyses further indicate that *Klk* genes are specifically expressed in granular convoluted tubule (GCT) and acinar cells [42], which are approximately 5-and 1.5-fold more abundant in males, respectively [27]. This raises the question of whether differences in cell abundance between male and female submandibular glands alone can explain the observed sex-biased expression. If expression per cell were equivalent between sexes, differences in cell-type abundance could, in principle, account for higher *Klk* expression in males. However, when we quantified overall expression levels, we found that male-female differences exceeded by at least 5.5-fold for all differentially expressed *Klk*s, indicating that the observed expression differences cannot be explained by cell-type proportions alone (**Figure 3c**).

One remarkable exception is that the observed male-biased pattern for *Klk1b* genes is not observed in *Klk1*. In fact, we found that *Klk1* is expressed significantly higher in female submandibular glands despite the fact that *Klk1b* genes were shown to be duplicates of *Klk1* [43, 44], and that *Klk1* is also known to be expressed in acinar and GCT cells which are much more abundant in males. To verify this transcriptional level observation at the protein and cellular levels, we performed immunohistochemical staining of both male and female submandibular gland sections for the female-biased *Klk1* gene, one of the few *Klk* family members for which a robust antibody is available (**Figure 3d)**. Consistent with the gene expression pattern, protein abundance was significantly higher in female granular convoluted tubules (**Figure S8**), resulting in greater overall *Klk1* expression in females despite the higher abundance of these cells in males. This observation suggests evolution of complex regulatory mechanisms in the Klk locus, distinct for *Klk1* and its paralogs.

Together, these results indicate that differences in cellular composition alone are insufficient to explain sex-biased *Klk* expression and suggest additional genetic or epigenetic regulation that drive this dimorphism.

### Sex-specific *Klk* expression is shaped by hormonal regulation, and lineage-specific 3D genomic architecture

To investigate the genetic basis of male-biased expression at the *Klk* locus, we first generated a gene tree incorporating gene family expansion and contraction estimates inferred by CAFE 5 and obtained from Ensembl [45]. We found that placental mammals, including other rodents, harbor up to 15 *Klk* genes, whereas mice and rats harbor approximately 17-25, indicating a burst of duplications in the murid lineage (**Figure S9**). Specifically, there are 13 additional tandemly duplicated copies in mice compared to the human genome (**Figure 4a**), all duplicated from *Klk1* [43, 44] and with male-biased expression.

**Figure 4.**
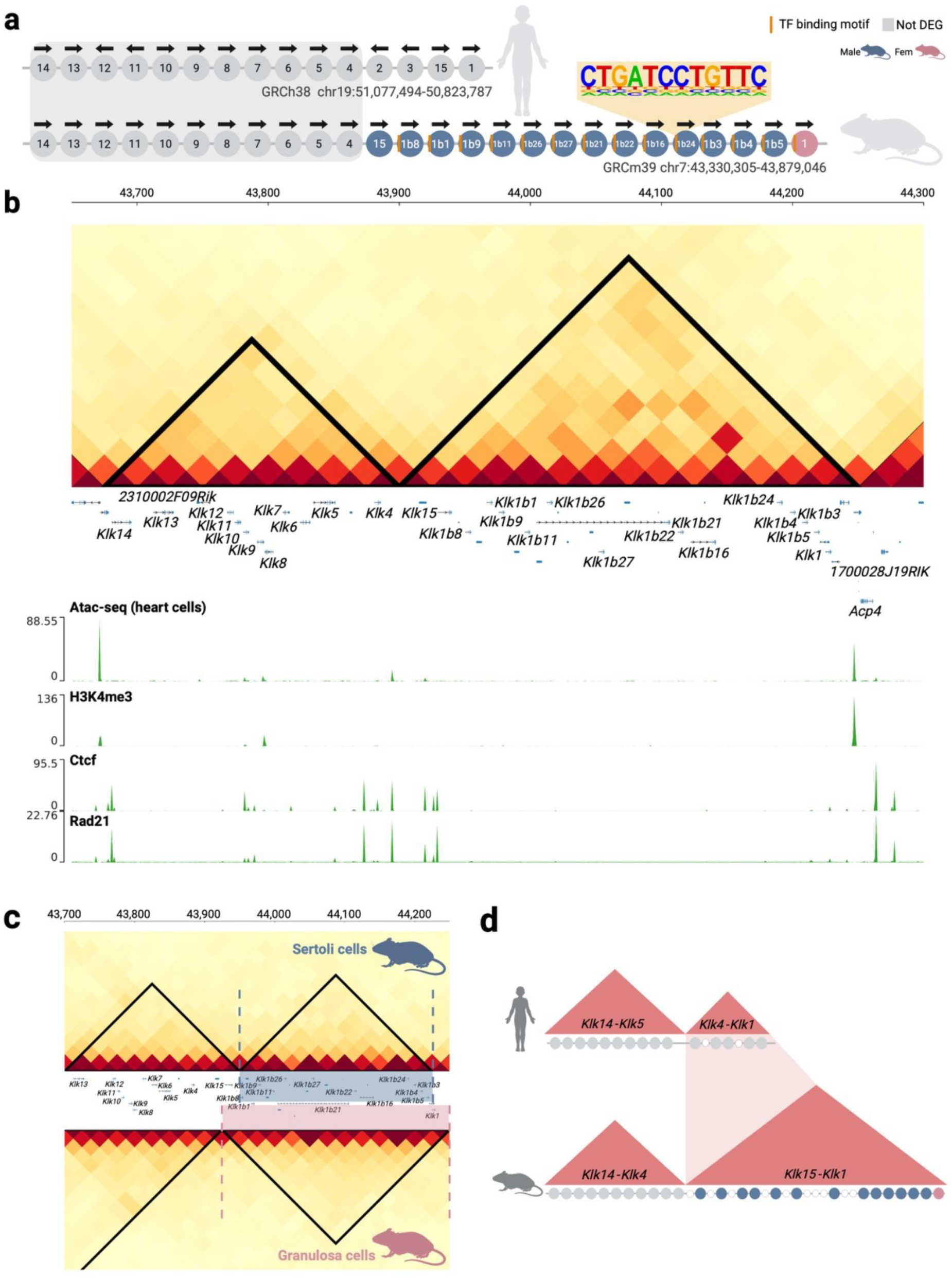
Expansion of the mouse *Klk* locus is associated with sex-biased expression and reorganization of local 3D genome structure. **(a)** Schematic comparison of the *KLK/Klk* locus organization in humans (top) and mice (bottom). Circles represent individual *KLK/Klk* genes ordered by genomic position and transcriptional orientation. Genes showing female-or male-biased expression are indicated in pink and blue, respectively. Orange bars denote putative transcription factor binding motifs shared across the expanded mouse cluster. The mouse lineage shows extensive tandem duplication of *Klk* genes compared with the more compact human locus. **(b)** Hi-C contact map of the mouse *Klk* region generated from B-cell lymphoma cell line, revealing two adjacent topologically associating domains (TADs). The left TAD predominantly contains non-sex-biased genes, whereas the right TAD encompasses the expanded cluster of sex-biased *Klk* paralogs. Gene annotations are shown below the contact map together with chromatin accessibility and regulatory marks (ATAC-seq, H3K4me3, CTCF, and RAD21), highlighting active promoters and architectural features associated with TAD boundaries. **(c)** Hi-C contact maps of the mouse *Klk* locus reconstructed from male Sertoli cells (top) and female granulosa cells (bottom). Dashed vertical lines indicate shifts in TAD boundaries between the two cell types. Genes located within these alternative domain configurations are highlighted, suggesting that differences in higher-order chromatin organization may influence sex-specific regulation of subsets of *Klk* paralogs. **(d)** Conceptual model illustrating how tandem gene duplication at the *Klk* locus may have been accompanied by expansion and restructuring of the surrounding TAD landscape.

Next, we asked whether the large cluster of differentially expressed genes containing *Klk* genes in the submandibular gland share common regulatory sequences that could underlie their coordinated male-specific expression. To address this, we performed a *de novo* motif enrichment analysis on this cluster, identifying overrepresented sequence motifs located near genes (see **Methods**). We found an enrichment (p = 1 × 10^-42^) for a conserved CTGATCCTGTTC motif located within 30 bp of the transcription start site in all sex-biased *Klk* genes except *Klk15* (**Figure 4a**; see Methods; **Tables S14** and **S15**). Previous work showed that this motif is a binding site for Ghrl2, a developmental transcription factor whose activity is sex-biased and correlated with testosterone levels [46, 47]. Consistent with this, kallikreins are well documented as androgen-regulated genes in the mouse submandibular gland [43, 48, 49]. In contrast, the motif is absent from *Klk* genes that do not show sex-biased expression. Moreover, none of the human *KLK* genes present this motif at the orthologous locus (**Table S15**). Given that the *Klk1b* paralogs arose through lineage-specific duplications of an ancestral *Klk1* gene [43, 44], our results support an evolutionary scenario in which the CTGATCCTGTTC motif emerged in the ancestral murine *Klk1* gene before subsequent gene expansion. Subsequent duplications may have increased gene expression through this regulatory mechanism, thereby amplifying male-biased expression in the submandibular gland.

Previous work in humans identified two distinct topologically associating domains (TADs) that structure the regulatory landscape of the *KLK* gene family, both demarcated by CTCF binding sites [50]. We therefore asked whether the lineage-specific duplications in mice resulted in the formation of a new TAD or expanded the existing one. To address this, we analyzed publicly available Hi-C and ChIP-seq datasets. Because salivary gland-derived Hi-C data were not publicly available, we used data from a mouse B-cell lymphoma cell line, which represented the highest-quality publicly available Hi-C dataset. Additionally, we used publicly available ATAC-seq data from mouse heart tissue to assess chromatin accessibility (See **Methods**). Consistent with the human architecture, we identified two TADs at the mouse *Klk* locus of which the second TAD is markedly expanded, encompassing 15 genes, including all those that show sex-biased expression (**Figure 4b**). Chromatin accessibility (ATAC-seq), histone modification (H3K4me3), and transcriptional anchor protein binding (Ctcf, Rad21) profiles delineate the borders of these TADs.

Although the TAD boundaries in mice largely mirror those observed in humans and encompass all sex-biased *Klk* genes, it remains unclear how *Klk1* retains both the regulatory motif and its position within the same TAD, while showing female-based expression. To explore whether sex-specific chromatin organization might contribute to this regulatory difference, we compared TAD boundaries between male and female cells. Using publicly available Hi-C data from male Sertoli cells and female granulosa cells, we reconstructed TAD structures and compared their boundaries (**Figure 4c**). This analysis revealed a slight boundary shift that coincides with the region showing male-specific expression, suggesting that *Klk1* in females may be regulated within a different TAD context. These findings indicate that regulatory control at this locus is more complex than previously appreciated, and that the data presented here are insufficient to fully explain the expression pattern of *Klk1*.

A major caveat of these analyses is that the cell types available for our analysis differ substantially from those of the submandibular gland in their developmental origin, transcriptional programs, and chromatin organization. These results should therefore be interpreted with caution until the observed TAD architecture is validated in salivary gland tissue and cell types. Regardless, they are consistent with the notion that lineage-specific Klk gene duplications have reshaped the ancestral TAD structure shared with humans (**Figure 4d**), expanding it in mice in a sex-specific manner, and offer an exciting framework for future investigation in relevant tissues and cell types.

## DISCUSSION AND CONCLUSION

In this study, we investigated the evolutionary forces shaping the composition of the salivary gland secretome in mice. We show that most highly expressed genes encoding secreted saliva proteins in mice are lineage-specific and lack direct human orthologs. We further identify the submandibular gland as the focal point of sex-specific expression. The kallikrein (*Klk*) gene family in mice accounts for ∼16.4% of male-biased expression, and we discuss how lineage-specific gene family expansion, emergence of novel regulatory motifs, and topological clustering contribute to the pronounced sex differences in expression. Together, the findings illustrate how gene duplication and regulatory rewiring drive sex-biased gene expression and contribute to the rapid evolution of the mammalian salivary gland secretome in different lineages.

When we examined sex differences in mouse saliva, we found that the submandibular gland is the main contributor to the protein differences observed between males and females, consistent with previous results [18, 28]. This male bias in submandibular expression is particularly notable when compared with the liver, the most studied tissue for sex differences in mice [30], where the degree of dimorphism is lower. We further found that this bias can be traced to the lineage-specific gene duplication clusters identified in our analysis, with the kallikrein (*Klk*) cluster standing out as a single gene family accounting for ∼16.4% of male-biased expression in the submandibular gland. The sex-biased expression of kallikrein genes is well documented in the literature, aligning with the patterns we observe here [38, 40, 41]. Specifically, our results support a model in which the expression of a hypothetical ancestral *Klk* gene in the murine lineage was subsequently amplified through a burst of tandem duplications within the same topologically associating domain (TAD), a process that could have both accompanied and facilitated developmental shifts in the male submandibular gland. This mechanistic framework suggests a broader evolutionary trend: rapid regulatory change across multiple genes can emerge through *de novo* motif evolution at a single locus, followed by its propagation via gene duplication within the same 3D genomic domain (**Figure 5**). A similar trend for motif evolution has recently been observed at the primate amylase locus [51].

**Figure 5.**
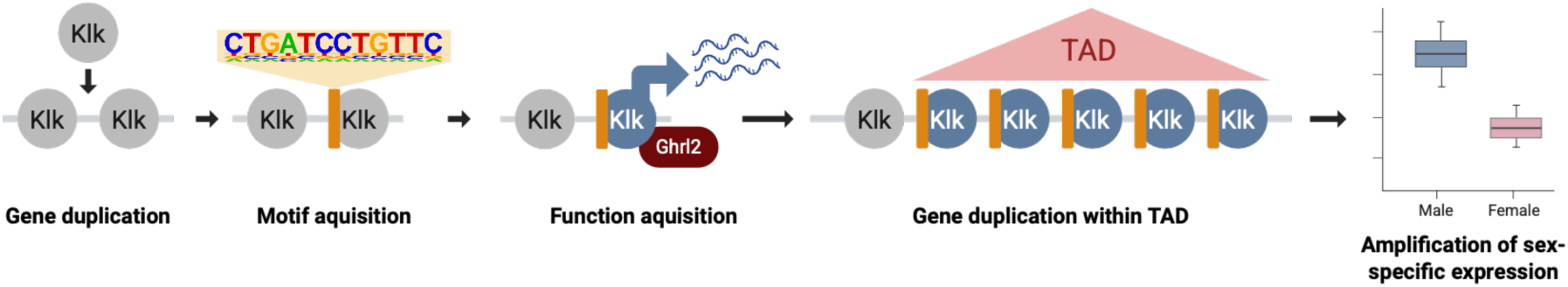
Proposed model for the evolutionary trajectory of mouse kallikrein (*Klk*) genes in salivary glands. Gene duplication created multiple *Klk* paralogs, which subsequently acquired new regulatory motifs. These regulatory changes enabled functional diversification, including interaction with Ghrl2. Further duplications occurred within a conserved topologically associating domain (TAD), promoting coordinated regulation of the expanded gene set. This combination of gene family expansion and regulatory rewiring led to increased male-biased expression in the submandibular gland.

The pronounced male-biased expression of kallikreins in saliva raises questions about their phenotypic impact in male mice. A prevailing hypothesis proposes that the presence of sex-specific proteins in saliva in male mice is related to improved healing through wound licking behavior after male-male aggression [38, 52]. While this hypothesis may be valid for the higher expression of wound healing related saliva proteins in male mice, such as epidermal growth factor and fibroblast growth factor [53, 54], it may not extend to the kallikrein gene family. An alternative hypothesis has been posited by Karn and Laukaitis where sex-biased expression of proteins in rodent saliva is explained by either signaling male dominance among rival males or mate identification through pheromone-like mechanisms [40].

Another fascinating parallel is that the syntenic region in reptiles, especially in snakes, underwent multiple, independent gene duplications and these lineage-specific kalikrein-like genes evolved to code for major venom proteins [55], also controlled by Ghrl-like transcription factor [56]. Similarly, a distant mammalian relative, shrew, evolved their venom through duplication of the Klk genes [57]. Further, earlier studies argued that Klk-like proteins from mouse saliva are toxic to non-mice rodents [58, 59]. Thus, within the context of male-male aggression, male-biased Klk genes may have evolved toxic properties. Our results cannot confirm or falsify either of these hypotheses, but they may provide a mechanistic base for formulating testable hypotheses approaching the questions from a molecular genetic level.

While we propose a hypothesis of co-regulation among male-biased *Klk* genes, our results do not fully explain the expression pattern of the only female-biased *Klk* gene (*Klk1*). Previous studies suggest that additional regulatory mechanisms contribute to the regulation of *Klk* transcripts, including microRNA-mediated silencing [60], highlighting the regulatory complexity of this locus. Moreover, *Klk1* has been reported to be regulated by estrogen in mice through mechanisms distinct from those described for other *Klk* family members [61]. Future studies integrating genomic, regulatory, and three-dimensional genome architecture data from salivary glands of the same individuals ideally using a single-cell resolution will be necessary to validate our proposed model of co-evolution among regulatory motifs, chromatin organization, gene duplications, and cell-type composition.

The biological functions and physiological relevance of the extensive species-and sex-specific differences in salivary gene and protein expression remain largely unknown. While addressing these functions was beyond the scope of the present study, our results provide a comprehensive catalogue of candidate genes and proteins that can now be used to test hypotheses about the roles of salivary gland-derived proteins in physiology, immunity, and behavior. One novel observation is the sex-specific glycosylation of the Muc19 protein in mouse saliva, suggesting that differences in glycosylation machinery, particularly the differential expression of sialyltransferases between males and females, contribute to sex-specific modification of secreted mucins. This stands in contrast to humans, where MUC19 expression is low in salivary glands and the protein activity is debated [37, 62]. Notably, genetic variation in human *MUC19* has been linked to complex introgression patterns in hominins and signatures of local selection in Indigenous American populations, marking this gene as a hotspot of adaptive evolution [63]. Together, our observations provide a foundation for future work aimed at dissecting the regulatory basis of species-specific expression, the molecular mechanisms underlying sex-specific glycosylation, and the evolutionary processes shaping *MUC19* variation across mammals.

Of importance for biomedical research, our findings raise more fundamental questions about the suitability of the mouse as a standard model for studying human salivary gland biology, a concern that has also been noted previously [64]. Focusing on salivary glands and their secreted products, we show that mice and humans differ fundamentally in both the expression patterns and composition of salivary proteins, including their post-translational modifications. With respect to disease relevance in humans, we found that several highly expressed Sjögren’s-related genes [42], including *Dcpp1*, *Scgb2b26*, and *Scgb2b27*, are mouse-specific and lack human counterparts, thus highlighting the importance of validating translational studies involving mice salivary glands and saliva in human models. In addition, our results underscore the importance of explicitly accounting for sex as a biological variable in studies of oral and salivary gland biology, particularly in mouse models.

Overall, this study, although limited to a comparison between two species, contributes to the emerging view that saliva exhibits considerable evolutionary plasticity, enabling accelerated evolution under selective pressures [4, 7, 9] and offers novel additional mechanistic hypotheses to explain these trends. Furthermore, our results are consistent with the hypothesis of developmental systems drift [65], whereby similar phenotypic traits across species evolve through distinct genetic or gene-regulatory mechanisms. Accumulating evidence indicates that developmental system drift may contribute to substantial transcriptomic divergence among species despite conservation of organismal phenotypes. For example, a recent study found extensive divergence in developmental gene expression between mouse and hamster lower molars despite relatively limited morphological divergence, with developmental trajectories diverging as much as those of the more morphologically divergent upper molars [66]. Notably, many regulatory changes were shared between upper and lower molars, suggesting concerted evolution of their developmental programs. The authors propose that evolutionary changes affecting one organ may also alter the development of related organs, even when their final morphology remains largely unchanged. A similar process could potentially contribute to the extensive transcriptomic divergence we observe between mouse and human salivary glands. Because regulatory programs are shared across salivary gland types, evolutionary changes affecting one gland or shared salivary functions could have pleiotropic consequences across glands, allowing substantial divergence in gene expression without correspondingly large changes in organ identity or function.

Saliva performs broadly conserved functions across mammals, primarily related to digestion, protection of oral tissues, microbial homeostasis, and pathogen defense, our results suggest that these functions may be achieved through different genes and varied regulatory architectures across species. In this context, the murine *Klk* locus provides a useful model for investigating how gene duplication, regulatory motif evolution, and 3D genome organization can shape the evolution of sex-biased gene regulation and the emergence of novel sexually dimorphic traits. Expanding similar analyses across multiple mammalian species will be crucial for uncovering general principles governing the evolution of salivary gland gene regulation and function.

## METHODS

### Sample collection for RNA-seq

A total of 12 mice (3 months old) were used, consisting of three biological replicates for each strain-sex combination: 3 female CD1, 3 male CD1, 3 female C57BL/6, and 3 male C57BL/6 (**Figure S10**). CD1 and C57BL/6 mice were selected as representative outbred and inbred strains, respectively, as both are widely used in laboratory studies of salivary glands [19–21]. Mouse samples were provided by UCSF. Animals were euthanized following the approved protocol AN201280-00I.

Euthanasia was performed using gradual CO_2_ inhalation followed by cervical dislocation as a secondary physical method to ensure death, as approved by the UCSF IACUC and in accordance with the 2020 AVMA guidelines. Animals were placed in 5L cages and 100% CO_2_ was introduced at a flow rate of 2L/min to induce loss of consciousness prior to respiratory arrest. Death was confirmed via the absence of heartbeat and pedal reflex before performing cervical dislocation.

From each mouse, we collected five glands: the three major salivary glands (parotid, sublingual, and submandibular glands), as well as the pancreas and liver, which were used as reference tissues.

### RNA-sequencing and analysis of mouse transcriptomes and differential expression analysis

Total RNA was extracted from each tissue, and RNA-seq libraries were prepared by Azenta Life Sciences (GENEWIZ) using their Standard/Strand-Specific RNA-Seq service with poly(A) selection for mRNA enrichment, followed by sequencing on an Illumina® NovaSeq™ platform (2×150 bp). Raw reads were processed with Trim Galore [67], which uses cutadapt for adapter trimming and FastQC for quality assessment, to remove low-quality bases and adapter sequences. Cleaned reads were then aligned to the *Mus musculus* reference genome (GRCm39) using HISAT2 [68], and gene-level quantification was performed with featureCounts [69], which assigns aligned reads to annotated genes and produces a matrix of raw read counts per gene. To avoid multimapping issues stemming from highly similar paralogous sequences, our featureCounts command was configured to exclude multimapping reads, so only uniquely mapping reads were included when quantifying gene expression. The specific parameters used in our pipeline are available in our GitHub repository.

Six parotid samples (two male and one female CD1 and three female C57BL/6) exhibited high expression of genes associated with muscle function and contraction, which is unusual in parotid gland transcriptomes. These samples were excluded from downstream analyses, as their expression profiles suggested possible contamination during tissue dissection. Next, we measured differential gene expression between sexes for each tissue using DESeq2 [70], applying a significance threshold of *p_adj_*< 0.05 and |log2 fold change| > 1. DESeq2 models raw read counts for each gene and compares expression levels between sexes, estimating differential expression while accounting for biological variability. Counts are internally normalized by library size and sequencing depth to allow accurate comparisons across samples, and significantly differentially expressed genes (DEGs) were identified for each tissue.

To assess sample quality and confirm proper clustering by biological variables, we performed principal component analysis (PCA) across all glands and separately for the three major salivary glands (**Figure S1**). For this analysis, the data were processed using standard DESeq2 pipeline with an analysis design that accounted for gland, strain, and sex. Before normalization and transformation through this pipeline, we removed genes with fewer than 10 total read counts across all samples to reduce noise from very lowly expressed transcripts. Therefore, our input files included 29,408 genes for the PCA of all glands (for Fig. S1a) and 28,025 genes for the PCA of salivary glands only (for Fig. S1b). This particular filtering strategy was based on previous work and is the standard practice for comparative transcriptomics [71].

As an additional validation, we reanalyzed the data using an independent pipeline based on fastp [72], Salmon [23], tximport/tx2gene [73], and DESeq2 [71]. Salmon estimates transcript abundance using a probabilistic model that accounts for reads compatible with multiple transcripts, rather than simply assigning each multimapping read to a single gene. Using this pipeline, we validated both the expression abundance analysis (**Table S14**) and the sex-biased differentially expressed genes (**Table S15**). Overall, the results were consistent with our original analyses. One difference that stands out is that Klk1, which was marginally female-biased in our original analysis, was not significantly female-biased in the Salmon-based analysis.

We then tested whether any gland had significantly more DEGs than the others. To assess statistical significance in the number of differentially expressed genes, we conducted pairwise chi-square tests between each pair of tissues and corrected for multiple testing using the false discovery rate (FDR). The code for this analysis is available on GitHub.

### Orthology and Secreted Gene Identification

We first sought to determine for all genes in each tissue the human orthologs and which were mouse specific. For this, we used an orthology reference table provided by The Jackson Laboratory (JAX) (https://www.informatics.jax.org/downloads/reports/HMD_HumanPhenotype.rpt), which lists mouse-human orthologous genes. Using this table, we grouped genes based on the number of orthologous relationships between species. Specifically, genes were classified as one-to-one when a single mouse gene mapped to a single human gene; one-to-many when a gene in one species mapped to multiple genes in the other; and many-to-many when multiple genes in both species were linked through orthology relationships. Genes without any annotated orthologs in the other species were classified as mouse-or human-specific.

Orthological classification and gene annotation remain a major source of error and bias in comparative genomic analysis. Therefore, to ensure that the orthology classifications from this database were not biased, we performed a parallel analysis using the orthology database from Ensembl release 116. Ensembl infers orthologous relationships through its comparative genomics pipeline and provides downloadable classifications of orthologs between mouse and human. We repeated our orthology-based analyses and found broadly similar results (**Figure S2**). Based on our manual assessment of the most highly expressed genes, we found that the Jackson Laboratory database provides more reliable orthology assignments between human and mouse than the Ensembl database. For example, *Amy1*, which belongs to a well-known gene family with multiple, independently evolved paralogs in both mice and humans, was correctly annotated as a many-to-many ortholog by the Jackson Laboratory but was not represented at all in the Ensembl orthology database.

Next, we aimed to characterize the contribution of highly expressed genes to salivary secretion by identifying secreted proteins and building an expression profile for each gland. Transcript abundance was calculated in transcripts per million (TPM) using the standard formula [74]. TPM values were computed for all genes in the three major salivary glands (parotid, sublingual, and submandibular). All genes in each gland were annotated as secreted or non-secreted based on the UniProt mouse proteome (https://www.uniprot.org/proteomes/UP000000589) [75] and human from Human Protein Atlas proteinatlas.org [76]. For the mouse proteome, specific annotations for salivary proteins were not available; therefore, we classified all genes as secreted or non-secreted based on general proteome annotations from UniProt. Combined with the orthology annotations described above, these data were used to generate Figure 1. We identified the 30 most highly expressed genes that code for secreted proteins in each salivary gland and plotted their expression levels (in TPM) to visualize gland-specific expression patterns using packcircle (https://mbedward.r-universe.dev/packcircles).

### Analysis of human transcriptomes

For humans, we reanalyzed publicly available RNA-seq data from human salivary glands (parotid, submandibular, and sublingual) generated by Saitou et al. (2020). Raw sequencing reads were processed as described above for mouse, using the human reference genome (hg38). Expression levels were quantified as transcripts per million (TPM), and mean TPM values were calculated for each gland. For liver and pancreas tissues, transcript data in TPMs was downloaded from GTEx and filtered for the same genes present in our dataset for salivary glands. TPMs were then recalculated for only those genes, and mean TPMs were calculated using all individuals. For orthology classification, please see the description of orthology classification above. Differential expression analysis was performed using DESeq2 [59] using raw gene counts only for submandibular gland samples.

### Correlation analysis of human-mouse transcriptomes

To assess the similarity of gene expression patterns between mouse and human salivary glands, we performed cross-species correlations of gene expression (mean TPM values) for each gland, focusing on one-to-one orthologs for genes with TPM>2 in each tissue. Because genes encoding secreted proteins and non-secreted proteins are subject to distinct evolutionary pressures, and secreted proteins are foften expected to evolve and diverge more rapidly [77, 78], we focused our analysis on genes encoding secreted proteins, which constitute the primary functional components of saliva. For the mouse, we used the annotated transcriptomes generated in this study for all glands. For the human data, we used the transcriptomic profiles from Saitou et al. (2020) for the major salivary glands (parotid, submandibular, and sublingual) and from GTEx for the pancreas and liver.

Spearman correlation coefficients (ρ) were calculated for each comparison. Finally, we note that these analyses involve data from different cohorts and species, and are therefore subject to potential batch effects; as such, these results should be interpreted with caution. Because of these technical limitations, we avoided direct comparisons of human and mouse expression data as a primary analytical approach. Nevertheless, we identified several one-to-one orthologs that exhibit dozens-to hundreds-fold differences in expression between human and mouse salivary glands (**Table S3 and S5)**, differences that are unlikely to be explained solely by batch effects and therefore warrant further investigation.

To investigate whether the correlations observed in salivary glands differed from those in liver and pancreas, we compared Spearman correlation coefficients across tissues. For each pair of tissues, we performed paired bootstrap resampling of shared orthologs to estimate the difference in Spearman correlations, the corresponding 95% confidence interval, and a two-sided bootstrap *p*-value. *P*-values were adjusted for multiple comparisons using the Benjamini-Hochberg method. The code used for this analysis is available on the dedicated GitHub repository.

### Mouse saliva collection

Saliva samples were collected from 4-month-old male (n=5) and female (n=5) C57BL/6 mice. Mice were anesthetized with isoflurane and administered the sialogogue pilocarpine dissolved in sterile saline via intraperitoneal (i.p.) injection. Pilocarpine was delivered at a final dose of 4.53mg/kg (concentration 0.68mg/mL; injection volume 200uL per 30g body weight). Saliva collection commenced 3 minutes post-injection and continued for 10 minutes using a pipette placed in the oral cavity. Samples were immediately snap-frozen and stored at-80 degrees Celsius for subsequent protein analysis.

### Human saliva collection

Saliva samples were collected from humans (male and female, n=5 each) using the passive drooling technique as previously described [79].

### SDS-PAGE

The 8-16% gradient pre-cast Novex Tris-glycine gel (LifeTechnologies; catalog no. XP08162BOX) was used to study the protein profiling of saliva samples based on the molecular size of the protein. Meanwhile, the samples were prepared for gel electrophoresis by mixing with a 2X sample buffer followed by boiling at 100^0^C for 10 min. The samples (∼15 μg of protein/well) were loaded into the wells, and the gel was run on an electrophoresis system (XCell SureLock Mini-Cell, Invitrogen) at 170 V for approximately 40 minutes. Subsequently, the gel was prepared for gel staining.

### Coomassie blue-PAS (Periodic acid Schiff) Staining

Protein bands were first stained using the Coomassie blue staining procedure. In brief, the procedure included three steps: a) fixation using fixing solution for 15 min twice, b) staining in Coomassie stain blue (R-250) solution for 1 hr., and c) destaining for 30 min thrice. After the destaining step, the gel was stained subsequently using the PAS staining procedure as described [79]. Stained gels were digitized using the GE Healthcare ImageScanner III.

### Protein identification from excised gel bands by LC-MS/MS

Protein bands of interest were excised from Coomassie blue-stained SDS-PAGE gels corresponding to the ∼150 kDa (male) and ∼130 kDa (female) regions using a clean surgical blade and processed for protein identification by mass spectrometry. Gel slices were subjected to in-gel digestion with trypsin following standard protocols. Peptides were extracted and analyzed by liquid chromatography coupled to tandem mass spectrometry (LC-MS/MS). The resulting spectra were processed using Spectronaut (Biognosys AG, version 20.0) and searched against the UniProt *Mus musculus* protein database. Protein identifications were filtered using a false discovery rate (FDR) threshold of 1% at both the peptide and protein levels.

### Lectin Blotting

Proteins were separated by SDS-PAGE under reducing conditions using 8-16% gradient gels and subsequently transferred to nitrocellulose membranes. For lectin detection, membranes were blocked in Tris-buffered saline containing 0.1% Tween-20 (TBST) supplemented with 2% Polyvinyl alcohol (PVA) for 1 h at room temperature. Membranes were then incubated with biotinylated lectins diluted in blocking buffer. Peanut agglutinin (PNA) was used to detect exposed Galβ1–3GalNAc (T-antigen) O-glycan structures, while Maackia amurensis lectin II (MAL-II) was used to detect α2,3-linked sialic acid residues [80, 81]. Following lectin incubation, membranes were washed extensively with TBST and incubated with Alexa fluor-488-conjugated neutravidin. Lectin binding was detected by enhanced chemiluminescence and imaged with a digital imaging system.

### Protein digestion

Saliva specimens were processed following the E3 protocol described recently [82]. In brief, around 20 μL of crude saliva was first mixed with 100 μL SDS buffer that contained 5% (w/v) sodium dodecyl sulfate (SDS), 100 mM Tris-HCl solution (pH 8.0) and final concentration of 10 mM tris(2-carboxyethyl)phosphine (TCEP) and 40 mM 2-chloroacetamide (CAA), and then boiled at 95 ⁰C for 10 min. Next, the lysate was mixed with 4x volume of 80% acetonitrile (CAN) aqueous solution and transferred to E3filters (CDS Analytical, Oxford, PA). Following a centrifugation step, the E3filters were washed with 200 μL of 80% ACN aqueous solution three times. For protein digestion, 1 mg of trypsin (Promega, WI) in a 200 μL of 50 mM triethylammonium bicarbonate (TEAB) buffer was added to E3filters and incubated at 37 ⁰C for 16-18 h with gentle shaking (300 rpm). To elute peptides, the E3filters were first centrifuged at 3,000 rpm and then flushed with two elution buffers, 200 ml of 0.2% formic acid in water, and 200 ml of 0.2% formic acid in 50% ACN, respectively. The eluents were dried in SpeedVac, and then subjected to desalting cleanup using C18 based StageTips (CDS Analytical, Oxford, PA). Dried peptide samples were stored under - 80 ⁰C until further analysis.

### LC-MS/MS analysis

The LC-MS/MS analysis was performed using an UltiMate 3000 RSLCnano system coupled with an Orbitrap Eclipse mass spectrometer and a FAIMS Pro Interface (Thermo Scientific). The dried peptides were resuspended into 13 μl of LC buffer A (0.1% formic acid in water) and 10 μl of it was then loaded onto a trap column (PepMap100 C18, 300 μm × 2 mm, 5 μm; Thermo Scientific) followed by separation on an analytical column (PepMap100 C18, 50 cm × 75 μm i.d., 3 μm; Thermo Scientific). A linear LC gradient flowing at 250 nL/min was applied from 1% to 25% mobile phase B (0.1% formic acid in ACN) over 125 min, followed by an increase to 32% mobile phase B over 10 min. The column was washed with 95% mobile phase B for 5 min, followed by equilibration with mobile phase A for 15 min. For the ion source settings, the spray voltage was set to 1.9 kV, funnel RF level at 50%, and heated capillary temperature at 275°C. The MS data were acquired in Orbitrap at 120K resolution, followed by MS/MS acquisition in data-independent mode following a protocol described previously [83]. The MS scan range (m/z) was set to 380–985, maximum injection time was 246 ms, and normalized AGC target was 100%. For MS/MS analysis, the spectra were acquired with Orbitrap at resolution of 30K. The isolation mode was Quadrupole, the isolation window was 8 m/z, and the window overlap was 1 m/z. The collision energy was 30%, AGC Target was 400K, and normalized AGC target was 800%. For FAIMS analysis, a 3-CV experiment (−40|−55|−75) was applied.

### Proteome quantitation and data analysis

The MS data were processed using Spectronaut software (version 20.0), a library-free directDIA+ workflow, and the UniProt protein databases (17,141 sequences; reviewed only). Briefly, the settings for Pulsar and library generation include: Trypsin/P as specific enzyme; peptide length from 7 to 52 amino acids; allowing 2 missed cleavages; toggle N-terminal M turned on; Carbamidomethyl on C as fixed modification; Oxidation on M and Acetyl at protein N-terminus as variable modifications; FDRs at PSM, peptide and protein level all set to 0.01; Quantity MS level set to MS2, and cross-run normalization turned on.

Differential protein abundance between male and female samples was assessed using the built-in statistical framework in Spectronaut (version 20.0). The protein LFQ method was set to Automatic (MaxLFQ); Quantity MS level was set to MS2, and the Quantity type was Area. The Candidates output from Spectronaut was used for quantitation analysis. Proteins were considered significantly differentially abundant between sexes if they met the thresholds of |fold change| > 1.5 and q-value < 0.05.

### Transcriptome-Proteome Integration and Correlation Analysis

Proteomic data were filtered to retain only proteins with a single corresponding gene annotation to avoid ambiguity in gene-protein assignments. For each protein, mean intensity-based absolute quantification (iBAQ) values were calculated separately for males, females, and across all samples. For visualization and correlation analyses, expression values were transformed as log_10_(1 + mean iBAQ). To enable transcriptome-proteome comparisons, we first filtered the transcriptomic dataset to genes annotated as secreted proteins (see description above). Proteins whose corresponding genes were present in this filtered transcriptomic dataset were then retained, resulting in 467 matched gene-protein pairs. Spearman correlation analyses were then performed using these matched protein-transcript pairs. It is important to note that each salivary gland contributes only partially to the overall proteome composition of mixed whole saliva, the oral fluid analyzed in this study. Therefore, strong correlations between glandular transcriptomes and the whole-saliva proteome are not necessarily expected, given the distinct spectrum of proteins secreted by each gland.

Furthermore, several highly expressed salivary gland transcripts were not detected by mass spectrometry, consistent with known technical limitations of proteomic assays for certain salivary proteins [84]. Conversely, some proteins detected in saliva at high abundance, such as serum albumin, are not synthesized by salivary glands but instead enter whole saliva through plasma exudation in the oral cavity, as reported previously [4, 85].

### Immunohistochemistry analysis

Submandibular glands were manually dissected and immediately fixed in 4% paraformaldehyde (PFA) at 4°C overnight. Following fixation, tissues were cryoprotected using a graded sucrose approach (10% sucrose w/v followed by 25% sucrose w/v in PBS). Tissues were then embedded in optimal cutting temperature (OCT) compound (Fisher Scientific, Cat # 4585) and cryosectioned at a thickness of 16µM. Sections were permeabilized with 0.5% Triton X-100 in PBS for 20 minutes at room temperature. Following permeabilization, sections were blocked for 1 hour at room temperature in a buffer containing 10% donkey serum (Jackson ImmunoResearch, Cat# 017-000-121) and 5% BSA (Fisher BioReagents, Cat# BP9700100) in PBS.

Sections were incubated with primary antibodies, Kallikrein 1 polyclonal antibody (1:300; Invitrogen, Cat# PA5-115481) and Nerve Growth Factor (NGF) polyclonal antibody (1:300; Sigma-Aldrich, Cat# N6655), diluted in antibody diluent overnight at 4°C. Following washing with 0.05% Tween-20 in PBS, antibodies were detected using Cy2-conjugated secondary Fab fragment antibodies (1:300; The Jackson Laboratory) incubated for 2 hours at room temperature. Nuclei were counterstained with Hoechst 33342 (1:1000; Anaspec Inc. Cat# AS-83218). Slides were mounted using Fluoromount-G (SouthernBiotech, Cat# 01100-01) and imaged using a Zeiss LSM 900 confocal microscope with a 40 X oil immersion objective.

Quantification of KLK-1 and NGF expression was performed using Fiji software. Regions of interest (ROIs) were manually delineated around salivary gland ductal structures identified by distinct morphology. The Mean Fluorescence Intensity (MFI) was calculated for each ductal ROI. Results are reported as MFI (a.u.).

### Clustering of DEGs, and motif enrichment analysis

We hypothesized that co-regulated gene clusters contribute to the differences in gene expression observed between strains and sexes. To test this, we focused on the submandibular gland for sex differences, as this tissue showed the highest numbers of DEGs between sexes. Using bedtools, we grouped DEGs into genomic clusters defined by a 100 kb window, such as the following example:

`bedtools cluster-i SM_sex_DEGs.sorted.bed-d 100000 > SM_sex_DEGs.clustered_100kb.bed`

To evaluate whether DEGs were more clustered than expected by chance, we performed a permutation analysis. Specifically, we randomly sampled 10,000 subsets of 1615 genes (matching the observed DEG count) from the total set of expressed genes and applied the same clustering procedure to each subset. From these permutations, we built an expected distribution of the number of genes per cluster size and compared it to the observed data (**Table S12**).

Next, we focused on the kallikrein (Klk) gene cluster in the mouse submandibular gland. Promoter regions for genes within the cluster were extracted and analyzed for motif enrichment using HOMER (Hypergeometric Optimization of Motif EnRichment) [86]. HOMER performs de novo motif discovery by identifying sequence motifs that are significantly enriched in a target set of sequences relative to a background set using a hypergeometric framework, and compares discovered motifs to known transcription factor binding motifs in curated databases (including HOMER and JASPAR libraries). For each gene in the cluster, sequences spanning −400 bp upstream to +100 bp downstream of the transcription start site (TSS) were analyzed. De novo motif discovery was performed using motif lengths of 8, 10, and 12 bp (Table S14), according to the following code

`findMotifs.pl “$gene_list_in cluster” mouse “out_folder/” \`

`-start-400-end 100 \`

`-len 8,10,12 \`

`-p 4`

To further confirm the motifs identified in the enrichment analysis and evaluate whether they are present across other Klk genes in mouse and in their human orthologs, we searched for these motifs across all Klk gene sequences in both species. Gene sequences for all annotated Klk genes were downloaded from the GRCm39 mouse genome assembly and the corresponding human genome assembly in FASTA format. Extracted sequences included the full gene regions encompassing coding sequences as well as 5′ and 3′ untranslated regions (UTRs). The short motif sequence identified by HOMER (CTGATCCTGTTC) was queried against these gene sequences using BLASTN with the blastn-short task, which is optimized for short query sequences, according to the following example:

`blastn-task blastn-short-query $motif.fa-subject $genes.fa - outfmt 6-out out.blastn.out`

This approach allowed us to identify exact or near-exact matches of the motifs within Klk gene sequences (**Table S15**).

### Evolution of the *Klk* genes, chromatin conformation and Topological Associated Domains Visualization

To investigate the evolution of the Kallikrein genes and gene expansions-contractions, we used CAFE 5 [45]. To investigate whether the sex-biased expression of Kallikrein (*Klk*) genes could be explained by differences in genome architecture and transcriptional regulation, we analyzed publicly available epigenomic datasets from the ENCODE project. Specifically, we visualized the Klk locus (chr7:43650000-44300000, mm10) using Hi-C contact maps at 25kb resolution generated from B-cell lymphoma cell line (https://www.encodeproject.org/files/ENCFF076GYR/) using HiCExplorer [87]. We further identified topologically associating domains (TADs) from the Hi-C data and overlaid them onto the figure. The code used for TAD calling and visualization is provided below.

`hicFindTADs \`

`--matrix ENCFF651SGY.mcool::/resolutions/25000 \`

`--chromosomes chr7 \`

`--minDepth 75000 \`

`--maxDepth 250000 \`

`--step 50000 \`

`--correctForMultipleTesting fdr \`

`--thresholdComparisons 0.05 \`

`--outPrefix TADs_heart_chr7_25kb_Local WIN2=“chr7:43650000-44300000”`

`hicPlotTADs --tracks 25kbTADs_selec.ini --region $WIN2 \`

`--outFileName 25kbTADs_selec.svg --dpi 300`

To explore potential regulatory features underlying the observed expression patterns, we overlaid chromatin accessibility and ChIP-seq data obtained from ENCODE. ATAC-seq data from postnatal mouse heart tissue were obtained from experiment ENCSR451NAE, using the processed bigWig file ENCFF384BWM. We used heart-derived ATAC-seq data because no ATAC-seq dataset from the CH12.LX cell line was available in the ENCODE database.CTCF ChIP-seq data from the mouse CH12.LX B-cell lymphoma cell line were obtained from experiment ENCSR000ERM, using the processed bigWig file ENCFF255NVD. RAD21 ChIP-seq data from the same cell line were obtained from experiment ENCSR000ERK, using the processed bigWig file ENCFF021SZL. H3K4me3 ChIP-seq data were obtained from experiment ENCSR000CGK, using the processed bigWig file ENCFF012DBS.

To investigate sex-specific differences in chromatin organization at the kallikrein (Klk) locus, we analyzed publicly available Hi-C datasets generated from granulosa (female) and Sertoli (male) cells [88]. These datasets were obtained from the 4DN Data Portal (https://data.4dnucleome.org/) under accession 4DNEXDK2R8M5 and 4DNEX99DGLEF. Multi-resolution cooler (.mcool) files were downloaded for each dataset (Sertoli: 4DNFIB7RFFBB; granulosa: 4DNFIZ59STGB). These cell types were selected because they represent sex-specific gonadal cell lineages, providing a model to examine potential sex-dependent chromatin architecture. Hi-C contact maps were extracted at 25 kb resolution, and topologically associating domains (TADs) were independently identified for each dataset using HiCExplorer as described above. The resulting TAD boundaries were then compared between male and female datasets to assess differences in higher-order genome organization at the Klk locus.

## DECLARATIONS

## Abbreviations

ACN: Acetonitrile;
AGC: Automatic gain control;
ATAC-seq: Assay for transposase-accessible chromatin using sequencing;
AVMA: American Veterinary Medical Association;
BSA: Bovine serum albumin;
CAFE: Computational Analysis of gene Family Evolution;
CAA: 2-chloroacetamide;
ChIP-seq: Chromatin immunoprecipitation sequencing;
CO₂: Carbon dioxide;
CTCF: CCCTC-binding factor;
DEG: Differentially expressed gene;
FAIMS: Field asymmetric ion mobility spectrometry;
FDR: False discovery rate;
GEO: Gene Expression Omnibus;
GCT: Granular convoluted tubule;
GTEx: Genotype-Tissue Expression project;
Hi-C: High-throughput chromosome conformation capture;
HOMER: Hypergeometric Optimization of Motif EnRichment;
iBAQ: Intensity-based absolute quantification;
IACUC: Institutional Animal Care and Use Committee;
i.p.: Intraperitoneal;
JAX: The Jackson Laboratory;
KLK/Klk: Kallikrein;
LC-MS/MS: Liquid chromatography-tandem mass spectrometry;
LFQ: Label-free quantification;
MAL-II: Maackia amurensis lectin II;
MFI: Mean fluorescence intensity;
MS: Mass spectrometry;
MS/MS: Tandem mass spectrometry;
NGF: Nerve growth factor;
OCT: Optimal cutting temperature compound;
PAR: Parotid gland;
PAS: Periodic acid-Schiff;
PBS: Phosphate-buffered saline;
PCA: Principal component analysis;
PFA: Paraformaldehyde;
PNA: Peanut agglutinin;
PSM: Peptide-spectrum match;
PVA: Polyvinyl alcohol;
ROI: Region of interest;
RNA-seq: RNA sequencing;
SDS-PAGE: Sodium dodecyl sulfate-polyacrylamide gel electrophoresis;
SL: Sublingual gland;
SM: Submandibular gland;
TAD: Topologically associating domain;
TBST: Tris-buffered saline with Tween-20;
TCEP: Tris(2-carboxyethyl)phosphine;
TEAB: Triethylammonium bicarbonate;
TPM: Transcripts per million;
TSS: Transcription start site;
UTR: Untranslated region.

## Ethics approval and consent to participate

All animal procedures were performed in accordance with institutional guidelines and were approved by the University of California, San Francisco Institutional Animal Care and Use Committee (IACUC) under protocol AN201280-0I.

## Consent for publication

Not applicable.

## Availability of data and materials

RNA-seq data generated in this study have been deposited in the NCBI Gene Expression Omnibus (GEO) under accession number GSE328310. Human salivary gland RNA-seq data were obtained from Saitou *et al.* (2020) and are publicly available in GEO under accession number GSE143702. Proteomic data generated in this study have been deposited in MassIVE and are currently available for editorial and peer review through reviewer access credentials; these data will be made publicly available upon publication. The code used for data processing, statistical analyses, and figure generation in this study is publicly available on GitHub at https://github.com/luanelandau/Mouse_Transcriptome_SG.

## Authors’ contributions

LJBL conceived the study, performed all computational and statistical analyses, generated figures, and drafted the manuscript. SJ performed saliva collection and biochemical analyses, including gel electrophoresis, and contributed to figure generation. NG performed immunohistochemistry experiments and contributed to figure preparation. AS contributed to study conceptualization and data analysis. JMK provided domain expertise and contributed to manuscript review and editing. SK contributed to study conceptualization, provided domain expertise and biological samples, and reviewed and edited the manuscript. SR contributed to study conceptualization, provided domain expertise, supplied mouse and human saliva samples, contributed to proteomic analyses, and secured funding. OG conceived and supervised the study, contributed to data interpretation, and secured funding. LJBL, OG, and SR wrote the manuscript. All authors reviewed the manuscript and provided feedback.

## Acknowledgements

We thank Alber Aqil, Charikleia Karagerogiou, Kendra Scherr, Vincent Lynch, and Petar Pajic for helpful discussions and insights. We thank Yanbao Yu for assistance with proteomic analyses and mass spectrometry. We thank Azenta Life Sciences for assistance with RNA-seq library preparation and sequencing. This work utilized computational resources provided by the Center for Computational Research at the University at Buffalo.

