## Supplementary Figures for "Rapid evolution of mouse saliva through regulatory rewiring and gene expansions"

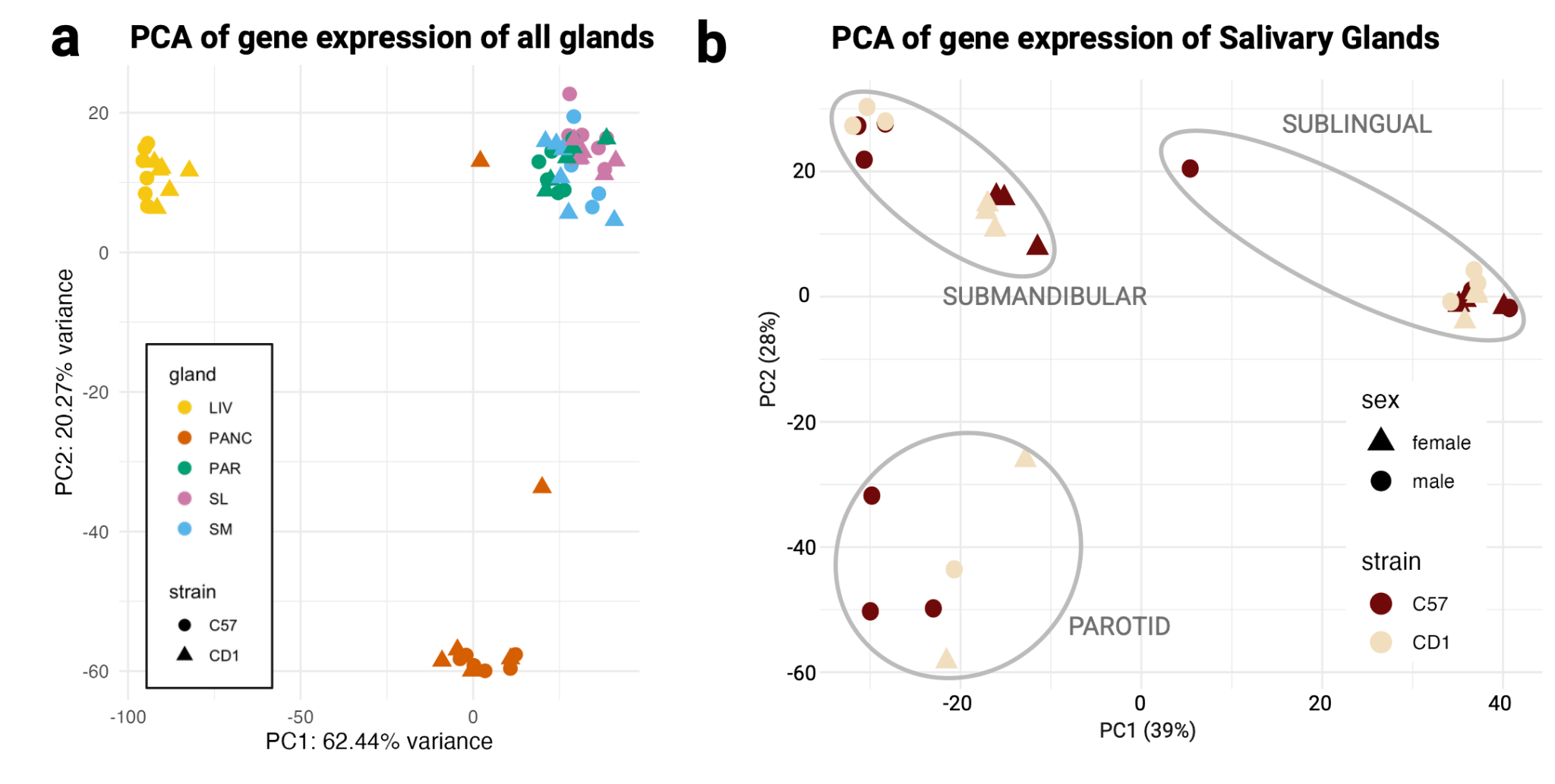

**Figure S1.** Samples cluster primarily by gland of origin. **(a)** Principal component analysis (PCA) of all samples. Colors indicate gland of origin (LIV=liver, PANC=pancreas, PAR=parotid, SL=sublingual, and SM=submandibular), and shapes indicate mouse strain (C57BL/6 and CD1). (b) PCA of salivary gland samples only. Colors indicate mouse strain (C57BL/6 and CD1), and shapes indicate sex (female and male).

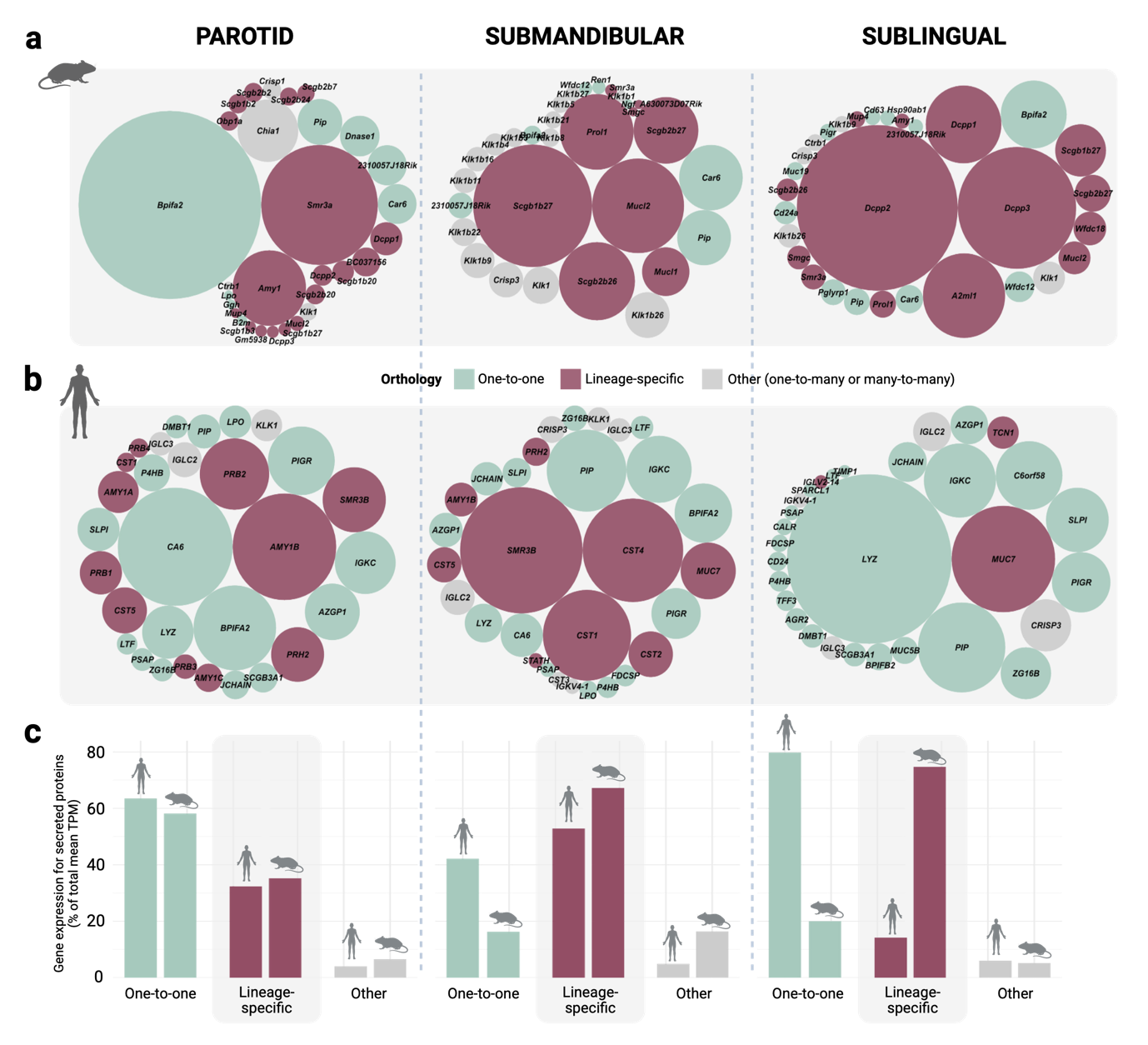

**Figure S2. Orthology classification of highly expressed genes coding for secreted proteins in mouse and human salivary glands using Ensembl orthology assignments.** (a) Bubble plots showing the top 30 most highly transcribed genes that code for secreted proteins in each major mouse salivary gland. Bubble size represents mean transcript abundance, estimated in transcripts per million (TPM), and colors denote orthology categories based on Ensembl assignments: one-to-one orthologs (green), lineage-specific genes (maroon), and other orthologous relationships (gray). (b) Same analysis as in (a), for human salivary glands. (c) Proportion of total gene expression (mean TPM) of genes coding for secreted proteins attributed to each orthology category in mouse and human.

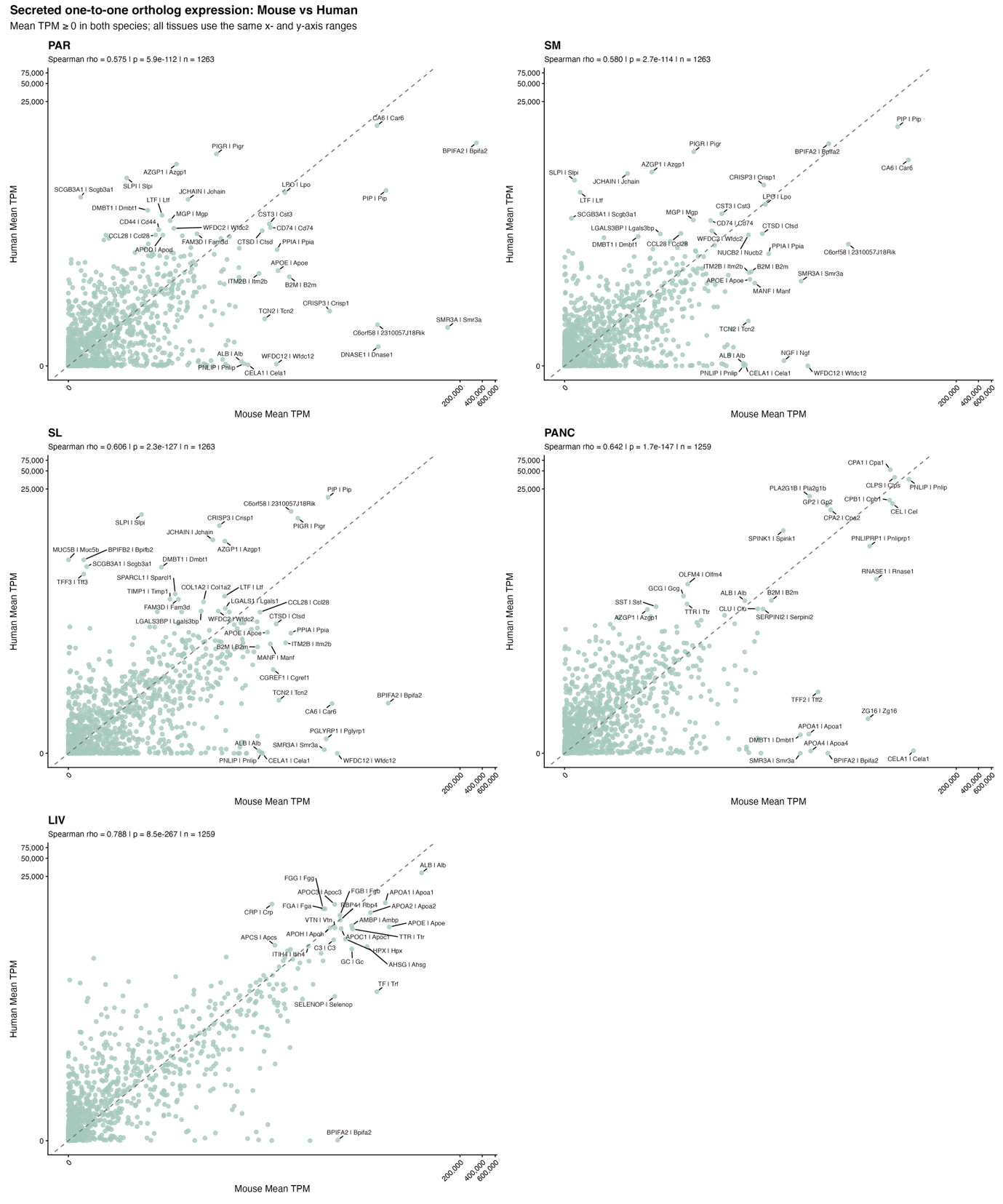

**Figure S3.** Cross-species comparison of secreted gene expression between mouse and human across tissues. Scatterplots show mean expression levels (log₁-transformed TPM) of one-to-one orthologous genes encoding secreted proteins in mouse (x-axis) and human (y-axis) for each tissue: parotid (PAR), submandibular (SM), sublingual (SL), liver (LIV), and pancreas (PANC). Each point represents a gene. The dashed line indicates equal expression between species (y = x). Spearman correlation coefficients are reported for each panel. Selected genes with high expression are labeled.

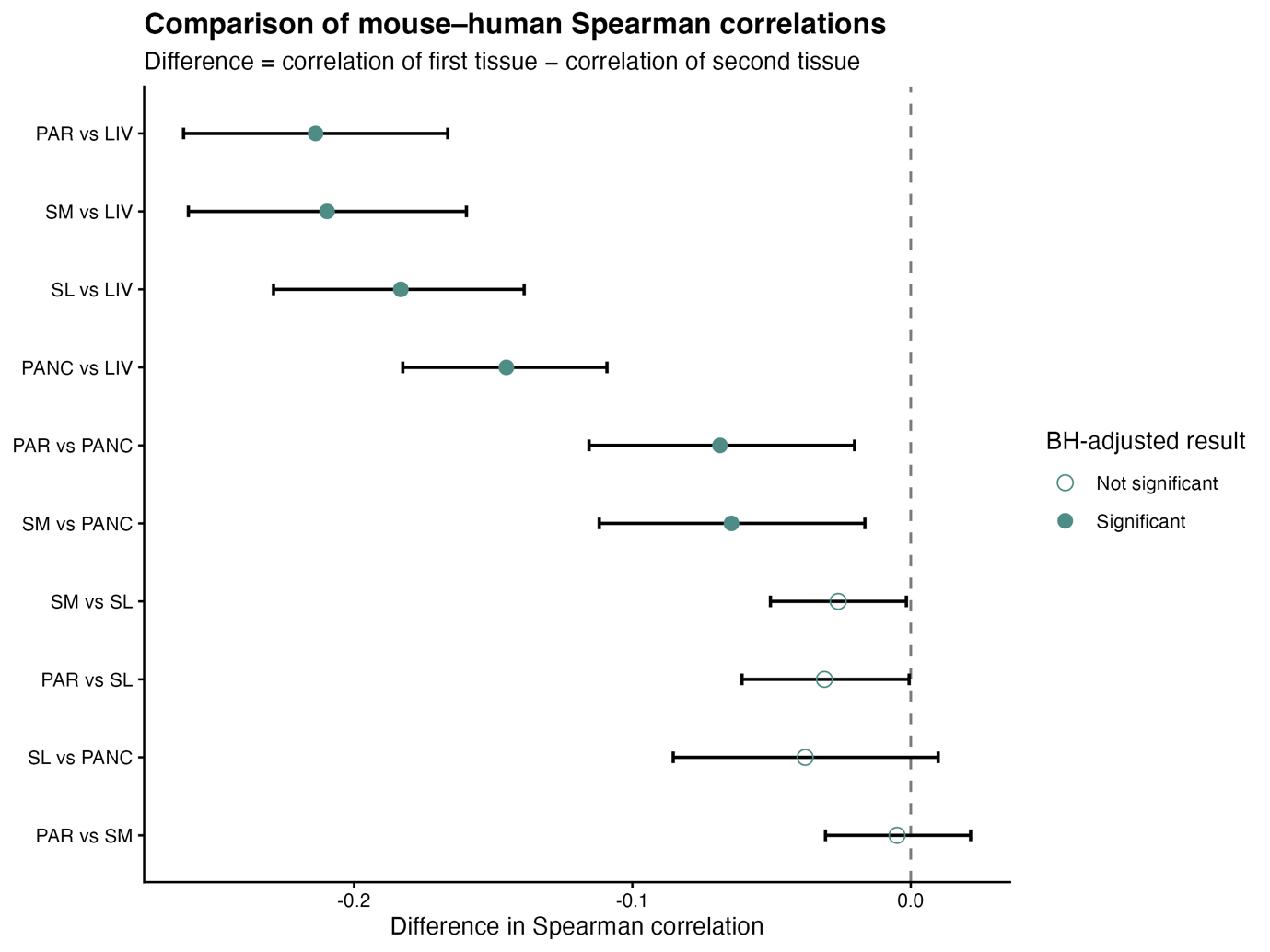

**Figure S4. Pairwise comparisons of mouse–human gene expression correlations across tissues.** Differences in Spearman correlation coefficients between tissues were calculated for the expression of one-to-one orthologous genes in mouse and human. Points represent the difference in Spearman correlation (correlation of the first tissue minus correlation of the second tissue), and error bars indicate 95% confidence intervals obtained by bootstrap resampling. The dashed vertical line indicates no difference between correlations. Filled points indicate comparisons significant after Benjamini-Hochberg (BH) correction for multiple testing (adjusted p < 0.05), whereas open points indicate non-significant comparisons. PAR, parotid gland; SM, submandibular gland; SL, sublingual gland; PANC, pancreas; LIV, liver.

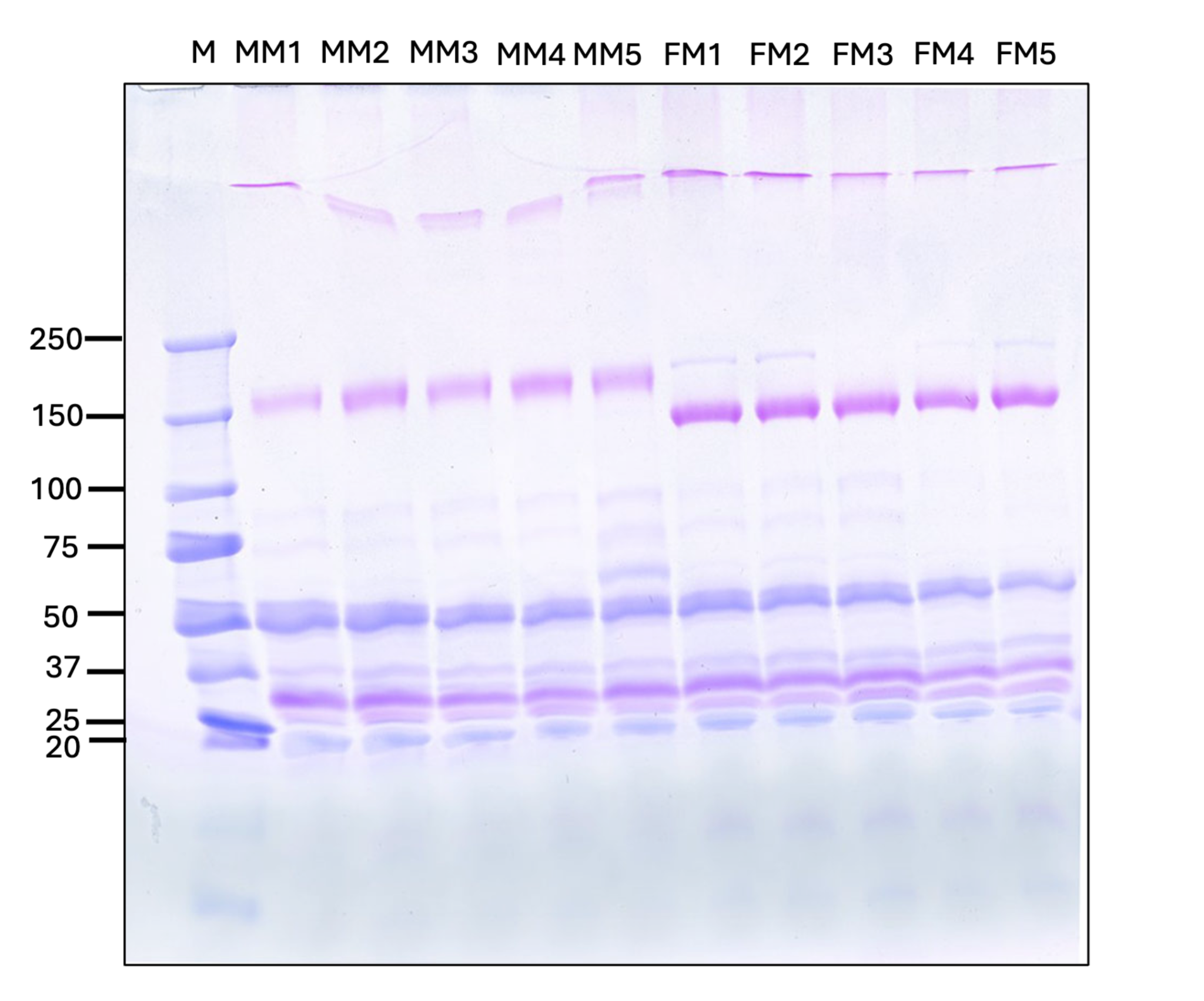

**Figure S5.** Sex-specific differences in mouse salivary protein profiles. Representative SDS-PAGE gel showing protein composition of whole saliva samples from male (MM) and female (FM) mice. M indicates the molecular weight marker. Each lane corresponds to an individual biological replicate. Differences in band intensity and migration patterns between males and females are visible across multiple molecular weight ranges, including prominent shifts in high-molecular-weight bands.

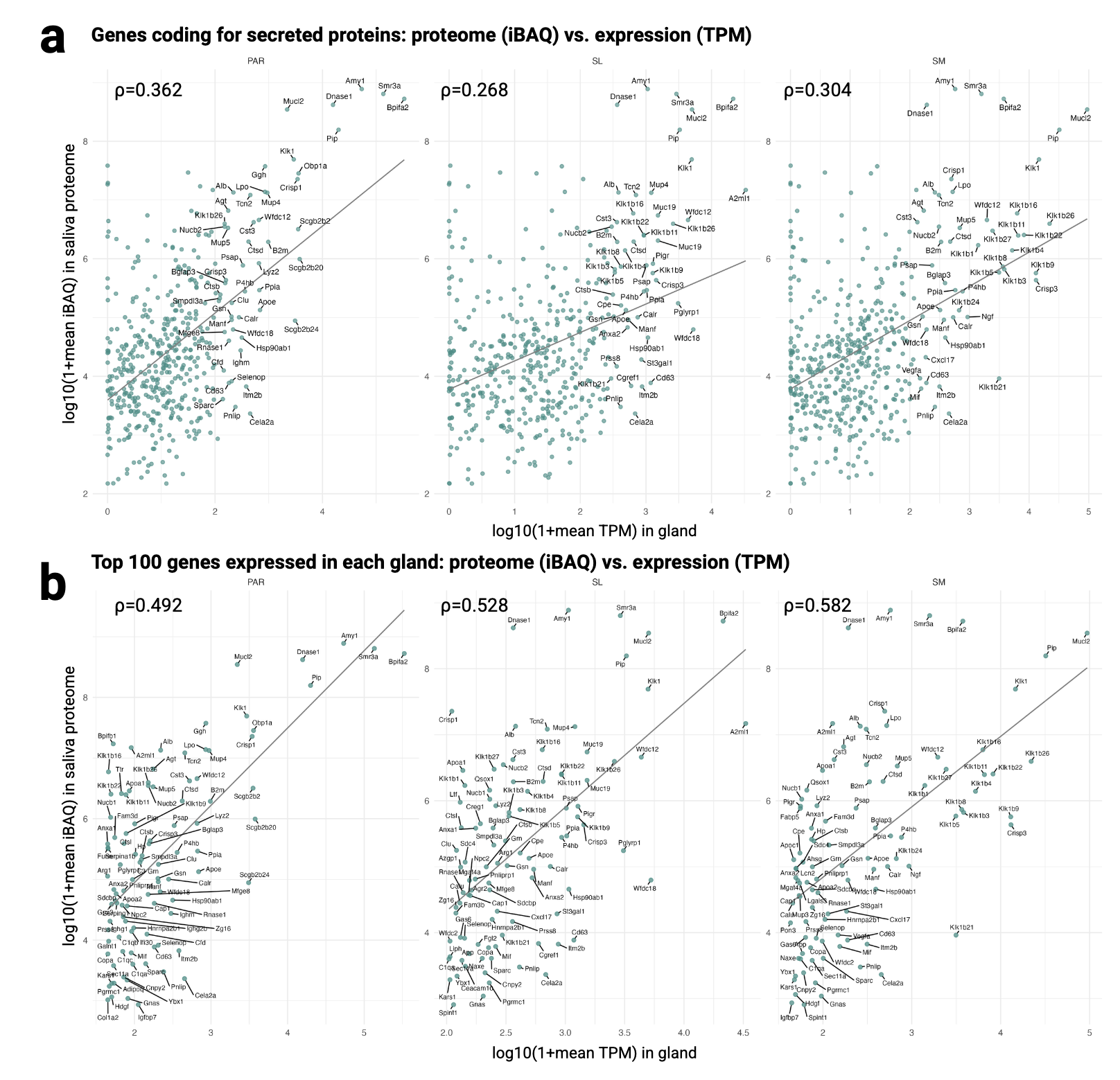

**Figure S6.** Correlation between transcriptomic and proteomic expression across salivary glands. **(a)** Scatterplots showing the relationship between gene expression (Log_10_(1+mean TPM) and corresponding protein abundance (Log₁₀(1+ mean iBAQ) for all matched gene-protein pairs in parotid (PAR), sublingual (SL), and submandibular (SM) glands. **(b)** Same as (a), but restricted to the top 100 most highly expressed genes encoding secreted proteins in each gland.

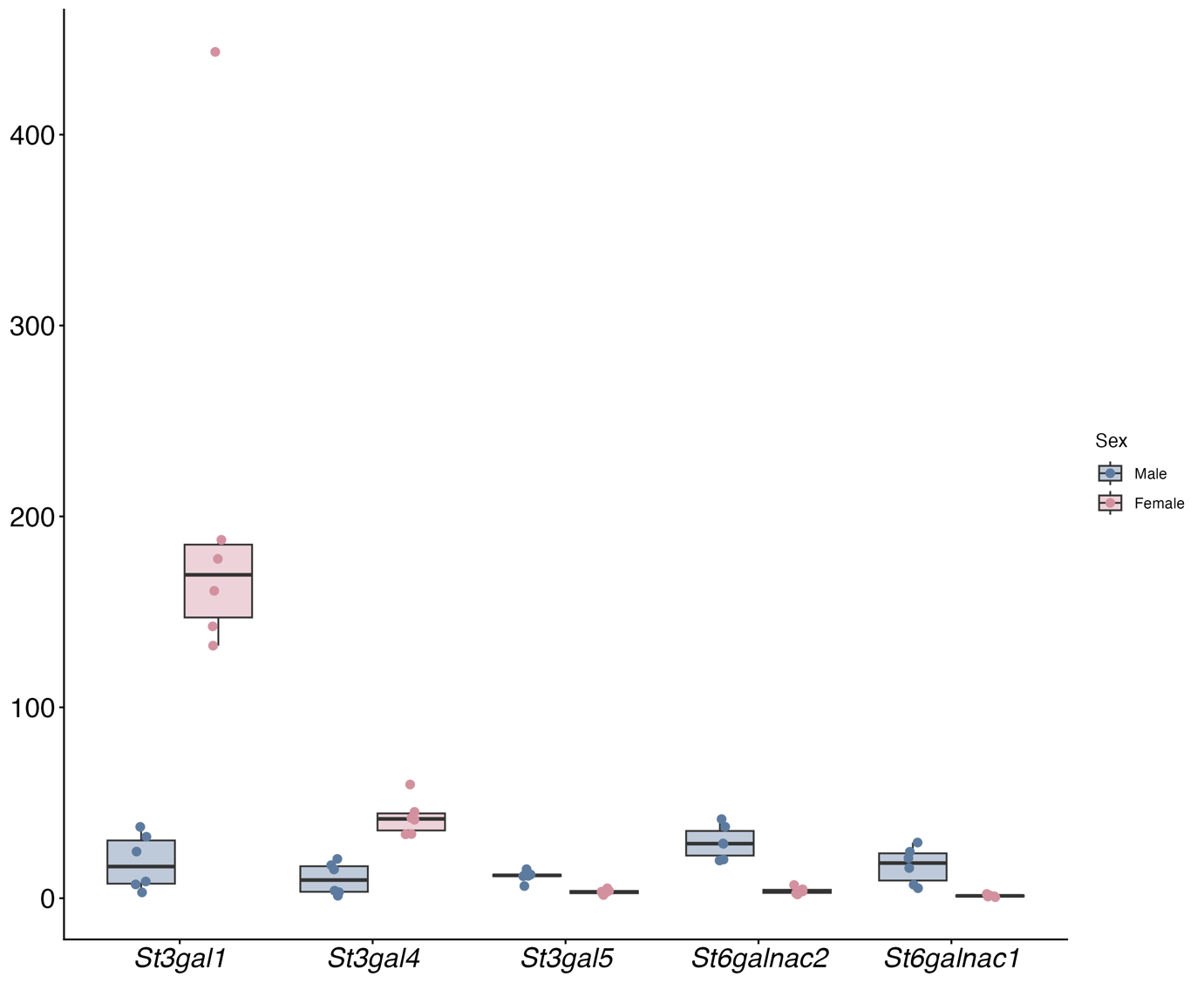

**Figure S7.** Sex-biased expression of sialyltransferase genes in the mouse submandibular gland. Boxplots show TPM expression levels of *St3gal1*, *St3gal4*, *St3gal5*, *St6galnac2*, and *St6galnac1* in male (blue) and female (pink) samples. Each point represents an individual sample. These patterns are consistent with sex-specific differences in protein glycosylation in mouse saliva.

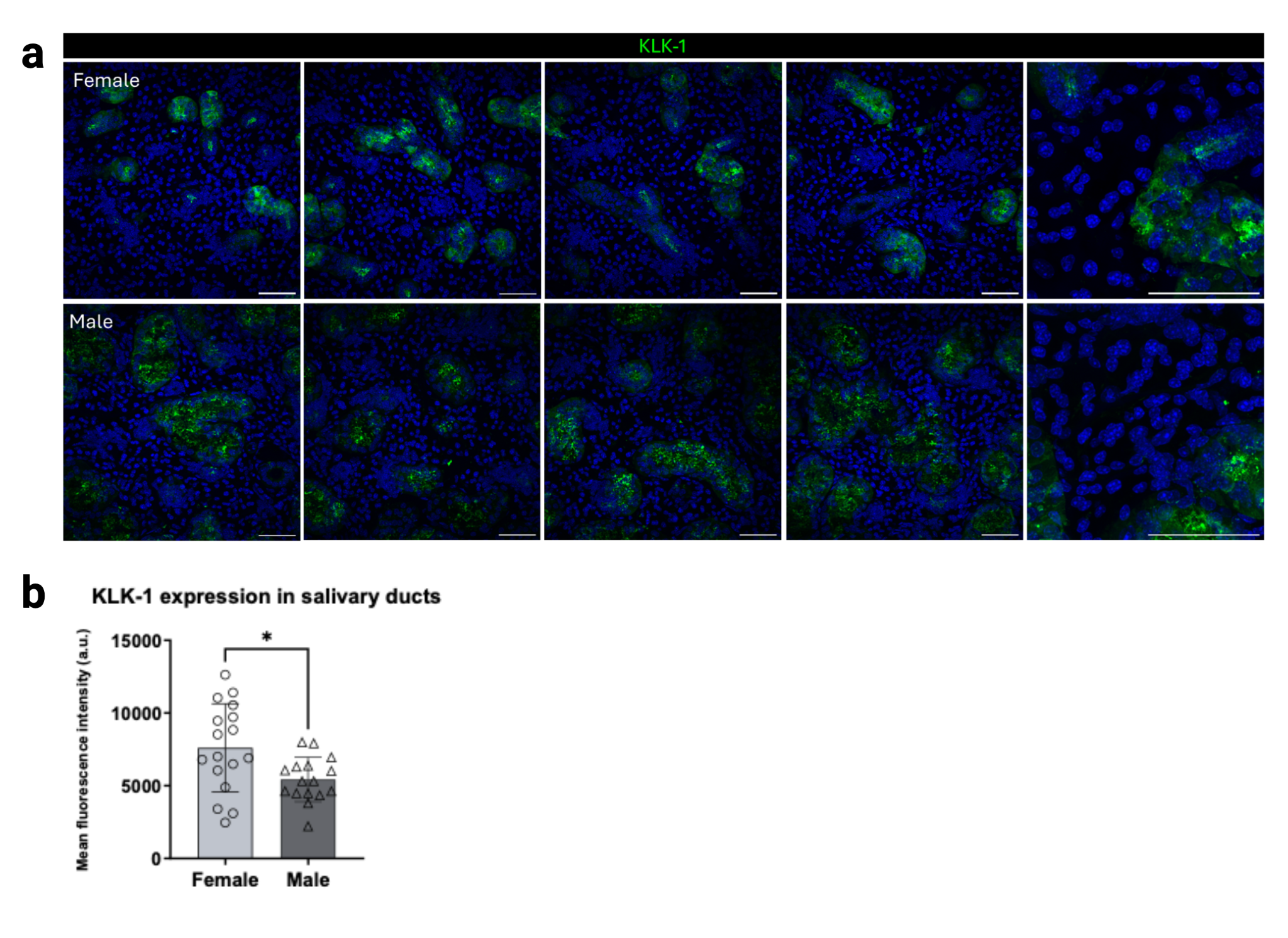

**Figure S8.** KLK1 shows higher abundance in female submandibular glands, despite the lower abundance of GCT cells in females.**(a)** Representative immunofluorescence images of KLK1 protein expression (green) in salivary gland tissue sections from female and male mice. Nuclei are stained with DAPI (blue). Multiple fields of view are shown for each sex. Scale bars are indicated. **(b)** Quantification of KLK1 fluorescence intensity across samples. Points represent individual biological replicates, and bars indicate mean ± standard deviation.

**Figure S9 is provided in PDF**

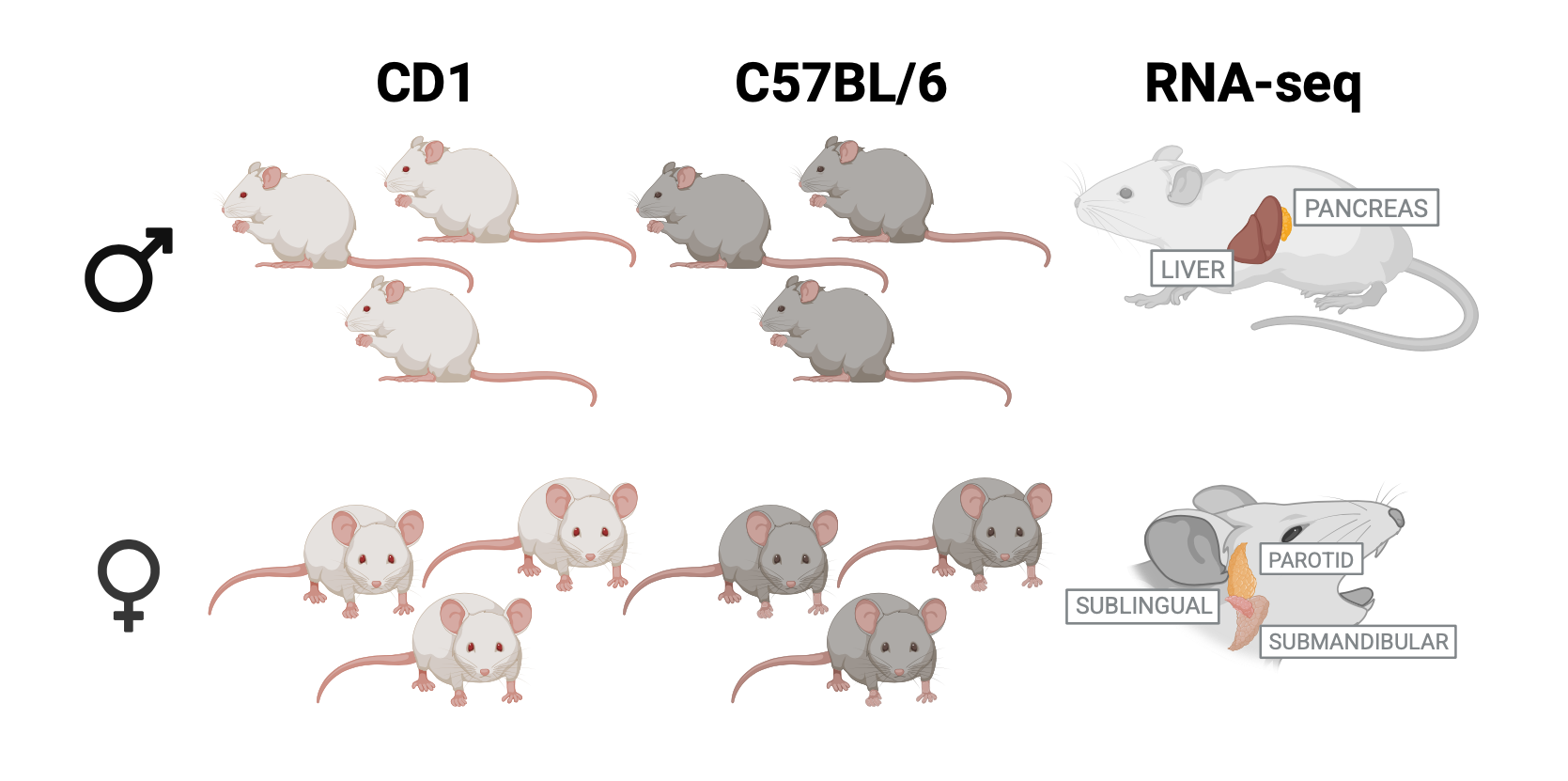

**Figure S10.** Overview of experimental design and tissue collection. Male and female mice from two strains (C57BL/6 and CD1) were used in this study. For each sex and strain, multiple biological replicates were collected. Tissues harvested included liver and pancreas, as well as the three major salivary glands: parotid, submandibular, and sublingual.
